# Attention field as a cognitive-behavioral marker for demarcating internet- but not smoking-addiction from reward

**DOI:** 10.1101/2024.12.23.630185

**Authors:** Mingxing Mao, Yaochun Cai, Ye Li, Zhuoqun Li, Wenshan Dong, Yuanyuan Wang, Xilin Zhang

## Abstract

Attentional effect (AE), attention profile (AP), and attention field (AF) have been studied extensively, however, their roles in addiction and demarcating addiction from rewards remain unclear. Using a modified Posner-paradigm with two types of pre-rewarded-cues (addiction-related and addiction-unrelated) and four groups (smoking-dependents, internet-dependents, and respective HCs), we found that both AEs and APs were independent of either cue type or group, while AFs were interactively modulated by the two. AFs of addiction-related cues were narrower than those of addiction-unrelated cues for internet-dependents, but not for either smoking-dependents or HCs; AFs of internet-dependents (not smoking-dependents) were narrower than those of HCs for addiction-related cues, but not for addiction-unrelated cues. Significantly, internet-dependents’ reduced AFs can be simulated by the divisive-normalization computation, both of which closely tracked their addictive severities. Our findings identify a cognitive-behavioral marker for demarcating internet-addiction from rewarding, arguing against the notion that internet-addiction, or, more generally, non-substance-addiction, is ill-posed.

## Introduction

Covert attention, the selective processing of visual information at a given location in the absence of eye movements, can be attracted automatically and involuntarily by salient stimuli, known as visual bottom-up attention [1–10]. The salience of stimuli is determined by not only purely physical properties but also various observer characteristics, such as for addiction patients, stimuli associated with addiction become extremely salient, which is not evident in individuals without a history of addiction. Indeed, atypical bottom-up attentional selection to addiction-related stimuli is a defining characteristic of various types of addiction, recently proposed as a potential diagnostic criterion. Specifically, addicted bottom-up attentional selection is marked by automatically and involuntarily allocating hypersensitive attention to addiction-related cues in the environment [11–16]. This bottom-up attentional bias for addiction-related cues is thought to result from acquired motivational and attention-grabbing properties of these cues due to sensitization of dopamine systems in the brain [17–21], and therefore, is increased during periods of subjective craving and has been shown to be predictive of treatment outcome and relapse in dependence [22–26]. Therefore, studying the addiction-related bottom-up attentional bias and abnormal attention processing will help improve the understanding of both the psychological and neurological mechanisms that manipulate attention selection, as well as their clinical remediation.

Intriguingly, although the addiction-related bottom-up attentional bias has consistently been found in various types of addiction, its clinical utility as a diagnostic criterion in DSM-V has not been well established [27, 28]. On the one hand, previous studies have argued that several of the hallmarks of this bias can be found in normal, healthy controls with no history of addiction following brief associative learning between arbitrary stimuli and reward outcomes [29–31], including the resistance to conflicting goals [29, 32–34], robustness to extinction [35] reliance on striatal dopamine signaling [36–38], and ability to facilitate approach behavior [39–41]. On the other hand, the addiction-related bottom-up attentional bias is evident in heavy but nondependent individuals [42, 43] and failures to predict later relapse have also been reported [44–46]. An important reason of this failure is that almost all previous studies investigated this topic focusing on the gain of attentional effect (AE) rather than its either field [7, 10, 47, 48] or profile [7, 49–53], a spatially scope and structured manner regarding how attention demarcates the target of interest from various distractors, respectively. This issue is particularly important since such attention profile (AP) and attention field (AF) are thought to closely reflect neural circuitry [48, 54–56], and therefore, could offer us a unique opportunity to give insight into the whole picture of addiction-related bottom-up attentional bias, with far-reaching implications for our understanding of social and neurobiological aspects underlying addiction. Consequently, to further establish the utility of addiction-related bottom-up attentional bias as a potential diagnostic criterion, it is very important to probe this bias free from such reward-driven attentional biases and examine both the APs and AFs of this attentional bias.

Here, we addressed these questions by using a modified Posner cueing paradigm with two types of pre-rewarded cues: addiction-related and addiction-unrelated, and manipulating the distance between the cue and probe to measure both the APs and AFs of addiction-related attentional bias for four groups: smoking-dependents, internet-dependents, and their respective demographic-matched healthy controls (HCs, Fig. 1), since that these two addictions are the predominant addictive behaviors in Chinese youth, which were the smoking addiction with the prevalence of nearly one third [57] and internet addiction which causing increasing detrimental effect on academic performance and mental health [58]. Results showed that both the cueing effects and their corresponding APs (monotonic gradient profiles) were independent of either the cue type or group, whose AFs, however, was interactively modulated by the cue type and group. The AF of addiction-related cues was significantly narrower than that of addiction-unrelated cues for internet-dependents, but not for either smoking-dependents or HCs. The AF of internet-dependents rather than smoking-dependents was significantly narrower than that of HCs for addiction-related cues, but not for addiction-unrelated cues. Significantly, the reduced AFs of internet-dependents can be simulated by the divisive-normalization computation of attention[48], both of which closely tracked internet-dependents’ addictive symptoms severities. Altogether, our findings reveal for the first time a unique AF alteration in internet-dependents, which can be considered as a cognitive-behavioral marker for demarcating internet-addiction, or, more generally, non-substance addiction, from arbitrary rewards.

**Fig. 1.**
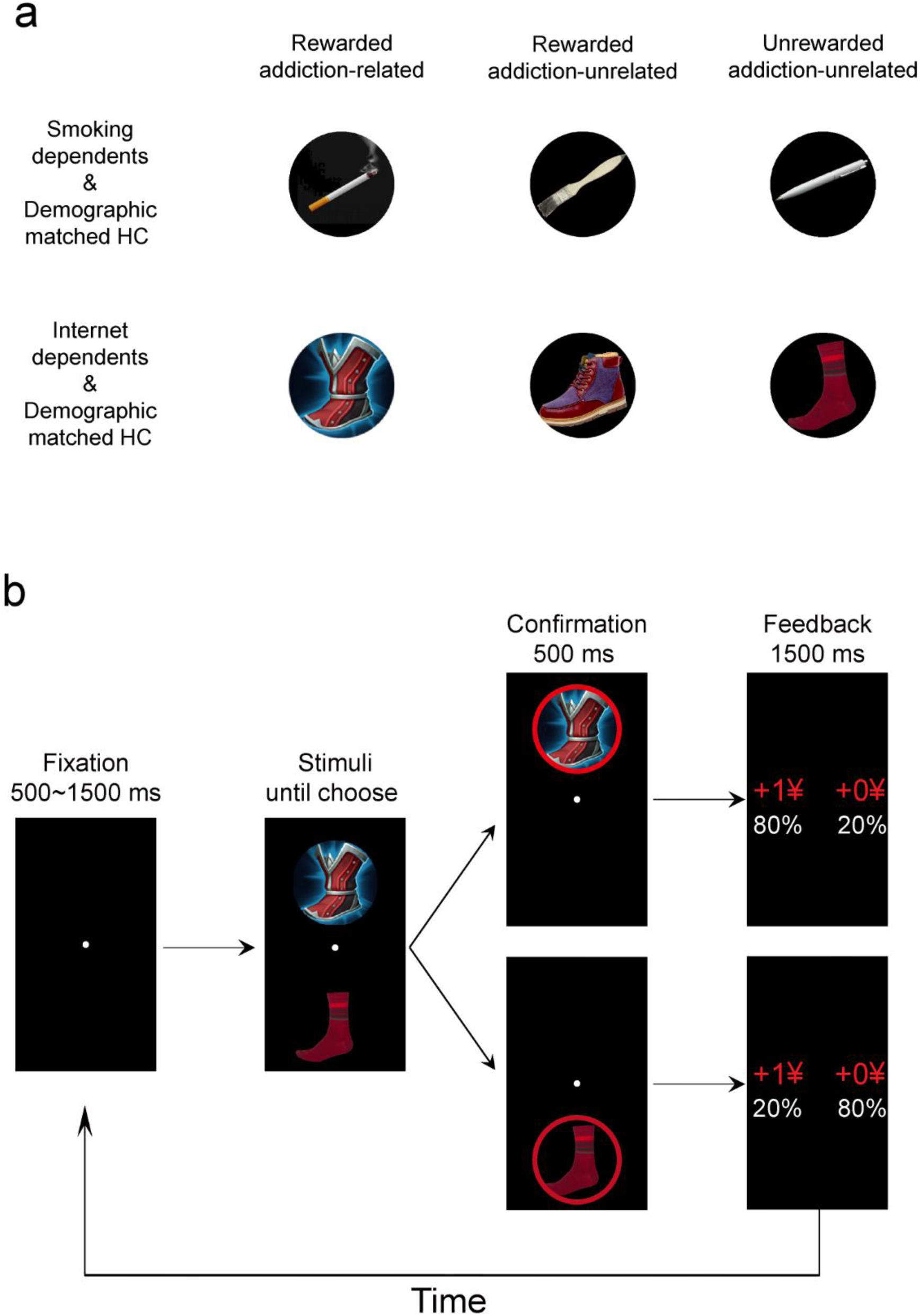
Stimuli and protocol in the training phase. a, For each dependent group and their demographic matched HCs, during the addiction-related reward condition, the probabilities of winning 1 ¥ were 0.8/0.2 for the addiction-related image and 0.2/0.8 for the addiction-unrelated image. During the addiction-unrelated reward condition, the probabilities of winning 1 ¥ were 0.8/0.2 for one of the addiction-unrelated images and 0.2/0.8 for the other. **b,** On each trial, one pair of stimuli was randomly presented and the relative position of two images was counterbalanced across trials. Subjects were required to choose either the upper or lower stimulus and their choice was circled in red. Then, the outcome was displayed on the screen, gain either 1 ¥ or 0 ¥. To win money, subjects had to learn, by trial and error, the stimulus–outcome associations. Subjects were told that they would play for real money, but at the end of the experiment their winnings were rounded up to a fixed amount.

## Results

Sample characteristics of 30 smoking-dependents, 30 demographic-matched HCs of smoking-dependents, 30 internet-dependents, and 30 demographic-matched HCs of internet-dependents, as well as their addiction-related measures are outlined in Table 1. Compared to their HCs counterparts, smoking-dependents reported significantly severer nicotine dependence (Fagerstrom test for nicotine dependence, FTND [59], t(29) = 10.558, *p* < 0.001, Cohen’s d = 3.921) and depressive symptoms (Beck Depression Inventory II [60], t(53) = 3.182, *p* = 0.002, Cohen’s d = 0.874), but not on gender, age, IQ as measured by Raven progressive matrices [61], or internet dependence measured by the Internet gaming disorder (IGD) [62] in Section 3 of DSM-V and Young’s Internet Addiction Test (YIAT) [63]. Compared to their HCs counterparts, Internet-dependents reported significantly more internet dependence on both the DSM-V IGD (t(58) = 12.446, *p* < 0.001, Cohen’s d = 3.268) and YIAT (t(45) = 8.843, *p* < 0.001, Cohen’s d = 2.636), but not on gender, age, depression, IQ or FTND.

**Table 1.** Demographic characteristics and addiction-related measures by study groups.

|  | Smoking-dependents (SDs) and HCs |  |  | Internet-dependents (IDs) and HCs |  |  |
| --- | --- | --- | --- | --- | --- | --- |
| | SDs (*n*=30) | HCs (*n*=30) | Test ($\chi^2$, *t*) | IDs (*n*=30) | HCs (*n*=30) | Test ($\chi^2$, *t*) |
| Gender (m/f) | 24/6 | 21/9 | *p* = 0.317 | 20/10 | 18/12 | *p* = 0.317 |
| Age (years) | 21.9 ± 0.36 | 20.90 ± 0.44 | *p* = 0.084 | 21.20 ± 0.40 | 20.27 ± 0.40 | *p* = 0.100 |
| Depression | **10.83 ± 1.45** | **5.13 ± 1.05** | ***p* = 0.002** | 9.93 ± 1.50 | 11.36 ± 1.64 | *p* = 0.522 |
| IQ | 49.33 ± 1.30 | 50.40 ± 1.25 | *p* = 0.556 | 50.13 ± 1.00 | 48.73 ± 1.24 | *p* = 0.384 |
| FTND | **4.03 ± 0.38** | **0 ± 0** | ***p* < 0.001** | 0 ± 0 | 0 ± 0 | *p* = 1.000 |
| DSM-5 IGD | 2.03 ± 0.40 | 1.37 ± 0.26 | *p* = 0.174 | **6.40 ± 0.27** | **1.40 ± 0.30** | ***p* < 0.001** |
| YIAT | 31.83 ± 3.75 | 24.10 ± 3.40 | *p* = 0.132 | **64.53 ± 2.22** | **23.13 ± 4.12** | ***p* < 0.001** |

*Depression: Beck Depression Inventory II; IQ: Raven progressive matrices; FTND: Fagerstrom test for nicotine dependence; IGD: Internet gaming disorder; YIAT: Young's Internet Addiction Test-gaming vision. $\chi^2$ tests were used for gender comparison; Independent samples *t*-test for between-group comparisons; Values are mean ± SE.*

### Training phase

In the training phase, all subjects performed a probabilistic instrumental learning task with monetary outcomes, adapted from previous studies [64–67]. Each of the pairs of stimuli was associated with pairs of outcomes (gain 1 ¥ or 0 ¥), and the two stimuli corresponding to reciprocal probabilities (0.8/0.2 and 0.2/0.8). For addiction-related reward conditions, the probabilities of winning 1 ¥ were 0.8/0.2 for the addiction-related image and 0.2/0.8 for the addiction-unrelated image. For addiction-unrelated reward conditions, the probabilities of winning 1 ¥ were 0.8/0.2 for one of the addiction-unrelated images and 0.2/0.8 for the other (Fig. 1a). On each trial, one pair was randomly presented and the relative position of two images was counterbalanced across trials. Subjects were required to choose either the upper or lower stimulus and their choices were circled in red. Then the outcome was displayed on the screen, gain either 1 ¥ or 0 ¥ (Fig. 1b). To win money, subjects had to learn the stimulus–outcome associations by trial and error. Subjects were instructed that they would play for real money, but at the end of the experiment their winnings were rounded up to a fixed amount.

We used a standard *Q*-learning algorithm [64, 68–70] to fit the reinforcement learning for each subject’s sequence of choices, and results showed that each condition demonstrated similar accuracy (*ACC*), learning rates (*α*) and exploration parameter (*β*) values (Fig. 2a and 2c). Each of these measurements was submitted to a two-factor mixed ANOVA with group (addictions and HCs) as the between-subjects factor and reward condition (addiction-related and addiction-unrelated) as the within-subjects factor. As in Fig. 2b, for smoking-dependents & HCs and each measurement, the main effect of group (*ACC*: F(1, 58) = 0.590, *p* = 0.446, partial eta squared, η_p_^2^ = 0.010; *α*: F(1, 58) = 0.152, *p* = 0.698, η_p_^2^ = 0.003; *β*: F(1, 58) = 0.298, *p* = 0.587, η_p_^2^ = 0.005), the main effect of reward condition (*ACC*: F(1, 58) = 0.120, *p* = 0.731, η_p_^2^ = 0.002; *α*: F(1, 58)= 0.391, *p* = 0.534, η_p_^2^ = 0.007; *β*: F(1, 58) = 0.057, *p* = 0.812, η_p_^2^ = 0.001), and the interaction between these two factors (*ACC*: F(1, 58) = 0.028, *p* = 0.868, η_p_^2^ < 0.001; *α*: F(1, 58)= 0.009, *p* = 0.927, η_p_^2^ < 0.001; *β*: F(1, 58) = 0.720, *p* = 0.400, η_p_^2^ = 0.012) were not significant. Similarly, as in Fig. 2d, for internet-dependents & HCs and each measurement, the main effect of group (*ACC*: F(1, 58) = 0.004, *p* = 0.951, η_p_ ^2^ < 0.001; *α*: F(1, 58) = 0.048, *p* = 0.828, η_p_^2^ = 0.001; *β*: F(1, 58) = 0.326, *p* = 0.570, η_p_^2^ = 0.006), the main effect of reward condition (*ACC*: F(1, 58) = 3.258 *p* = 0.076, η_p_^2^ = 0.053; *α*: F(1, 58) < 0.001, *p* = 0.994, η_p_ ^2^ < 0.001; *β*: F(1, 58) = 10.105, *p* = 0.002, η_p_ ^2^ = 0.148), and the interaction between these two factors (*ACC*: F(1, 58) = 0.374, *p* = 0.543, η_p_^2^ = 0.006; *α*: F(1, 58) = 0.574, *p* = 0.452, η_p_^2^ = 0.010; *β*: F(1, 58) = 0.171, *p* = 0.681, η_p_^2^ = 0.003) were not significant. These results strongly suggested that subjects in our training phase were rewarded equally well for each group and each condition.

**Fig. 2.**
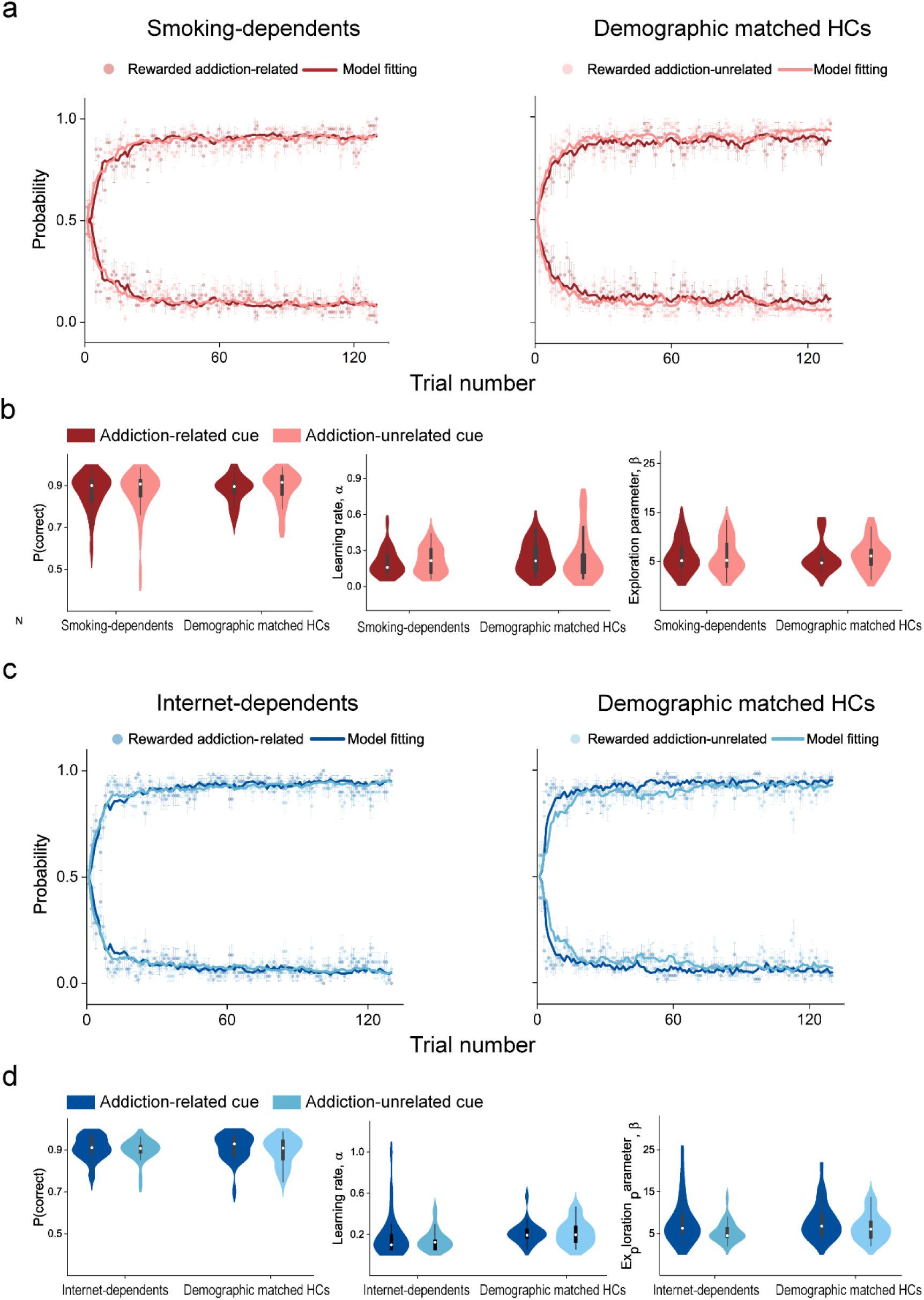
Results of the training phase. a, The dots depict the proportion of trials that subjects chose the ‘correct’ (associated with a probability of 0.8 of winning 1 ¥) and ‘incorrect’ (associated with a probability of 0.2 of winning 1 ¥) stimuli during addiction-related and addiction-unrelated reward conditions, for smoking-dependents (left) and their HCs (right). The curves represent the probabilities predicted by the computational model for each condition. **b**, The accuracy, learning rates (*α*), and exploration parameter (*β*) values for each group and each condition. **c** and **d**, Results of internet-dependents and their HCs, see caption for (**a** and **b**) for a description of each type of graph. Smoothed density plot denotes the distribution of the sample, box plot denotes the quantile and median scores, and error bars denote 1 SEM calculated across subjects.

### Test phase

Immediately following the training phase, subjects completed the test phase. The test phase consisted of two sessions (addiction-related and addiction-unrelated), which were the same except for the exogenous cue: addiction-related cue (ARC) or addiction-unrelated cue (AUC) (Fig. 3a). Subjects performed two sessions on the same day, and the order of the two session was counterbalanced across subjects. For both sessions, each trial began with the fixation. A cue frame with (the cue condition) or without (the non-cue condition) exogenous cue was equiprobably and randomly presented for 50 ms, followed by a 150 ms fixation interval. Then a probe grating, orientating at 45° or 135° away from the vertical, was presented for 50 ms. Subjects were asked to press one of two buttons as rapidly and correctly as possible to indicate the orientation of the probe (45° or 135°) (Fig. 3c). There was no significant difference in accuracy (Fig. S1) or removal (Fig. S2) rate (i.e., correct reaction times shorter than 200 ms and beyond 3 SDs from the mean reaction time in each condition were removed) across conditions (all p > 0.05). The cueing effect for each distance (D0-D4, Fig. 3b) was quantified as the difference between the reaction time (RT) of the probe task performance in the non-cue condition and that in the cue condition.

**Fig. 3.**
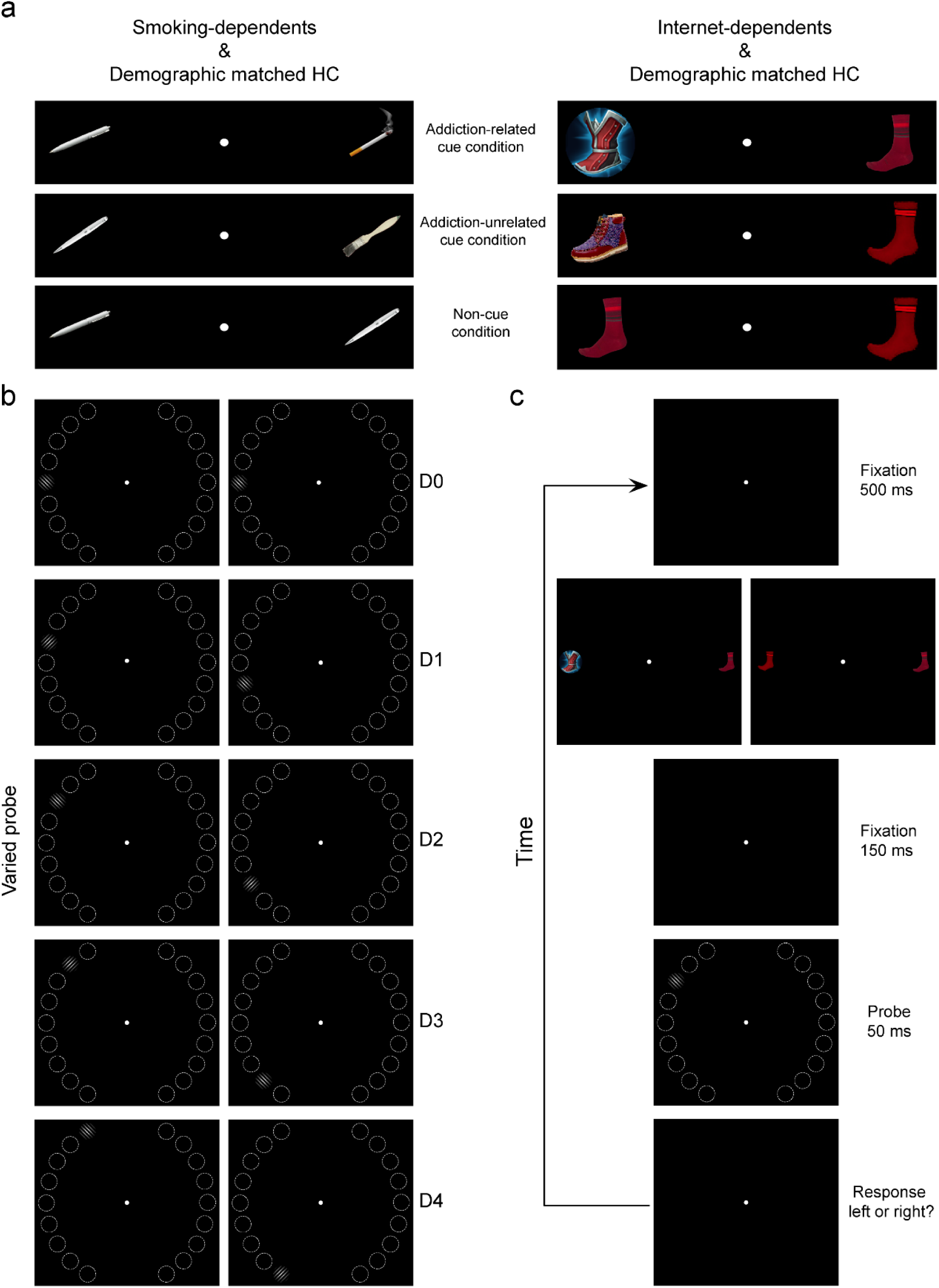
Stimuli and protocol in the test phase. a, For each dependent group and their demographic matched HCs, during the addiction-related cue condition, a pair of stimuli consisted of an addiction-related image and an addiction-unrelated image. The addiction-related image was rewarded in the training phase and served as an exogenous cue to attract subjects’ covert spatial attention. During the addiction-unrelated cue condition, a pair of stimuli consisted of two addiction-unrelated images, one of which was rewarded in the training phase and served as an exogenous cue. During the non-cue condition, a pair of stimuli consisted of two addiction-unrelated images, neither of them was rewarded in the training phase. For each condition, the relative position of two images was counterbalanced across trials. **b,** Each texture stimulus contained 18 positions, indicating by the dashed white circles (not displayed during the experiments), settled at an iso-eccentric distance from fixation; half of them were located in the left visual field, and the other half were located in the right visual field. In each visual field, the exogenous cue appeared (the cue condition, either addiction-related or addiction-unrelated) or was absent (the non-cue condition) in the middle of 9 positions with equal probability. The probe appeared equiprobably and randomly at 1 of the 9 possible positions and there were 5 possible distances between the cue and probe, ranging from D0 through D4. **c,** Psychophysical protocol. Each trial began with the fixation. A cue frame with (the cue condition) or without (the non-cue condition) exogenous cue was presented for 50 ms, followed by 150 ms fixation interval. Then a probe grating, orientating at 45° or 135° away from the vertical, was presented for 50 ms. Subjects were asked to press one of two buttons as rapidly and correctly as possible to indicate the orientation of the probe (45° or 135°).

### Attentional effect

Fig. 4 shows the cueing effect of each condition for both sessions; most of these cueing effects were significantly > 0, indicating that the bottom-up attention of the subject was attracted to the exogenous cue location, allowing them to perform more proficiently in the cue condition than the non-cue condition of the probe task. To examine whether the cueing effect can demarcate addiction (internet- and smoking-dependents) from arbitrary rewards, for each addiction group and their HCs, both the peak (i.e., D0) and mean of cueing effect were submitted to a two-factor mixed ANOVA with group (addictions and HCs) as the between-subjects factor and cue (ARC and AUC) as the within-subjects factor. For smoking-dependents and their HCs, the main effect of group (Peak: F(1, 58) = 0.285, *p* = 0.596, η_p_^2^ = 0.005; Mean: F(1, 58) = 0.230, *p* = 0.634, η_p_^2^ = 0.004), the main effect of cue (Peak: F(1, 58) = 1.588, *p* = 0.213, η_p_^2^ = 0.027; Mean: F(1, 58) = 1.990, *p* = 0.164, η_p_^2^ = 0.033), and the interaction between these two factors (Peak: F(1, 58) = 0.285, *p* = 0.596, η_p_^2^ = 0.005; Mean: F(1, 58) = 0.197, *p* = 0.659, η_p_^2^ = 0.003) were not significant (Fig. 4a). Similarly, for internet-dependents and their HCs, the main effect of group (Peak: F(1, 58) = 0.044, *p* = 0.834, η_p_^2^ = 0.001; Mean: F(1, 58) = 0.935, *p* = 0.338, η_p_^2^ = 0.0116), the main effect of cue (Peak: F(1, 58) = 1.801, *p* = 0.185, η_p_^2^ = 0.030; Mean: F(1, 58) = 0.750, *p* = 0.390, η_p_^2^ = 0.013), and the interaction between these two factors (Peak: F(1, 58) = 1.980, *p* = 0.165, η_p_^2^ = 0.033; Mean: F(1, 58) = 3.001, *p* = 0.089, η_p_^2^ = 0.049) were not significant (Fig. 4b). These results strongly suggested that the gain of attentional effect could not demarcate either internet-dependents or smoking-dependents from arbitrary rewards.

**Fig. 4.**
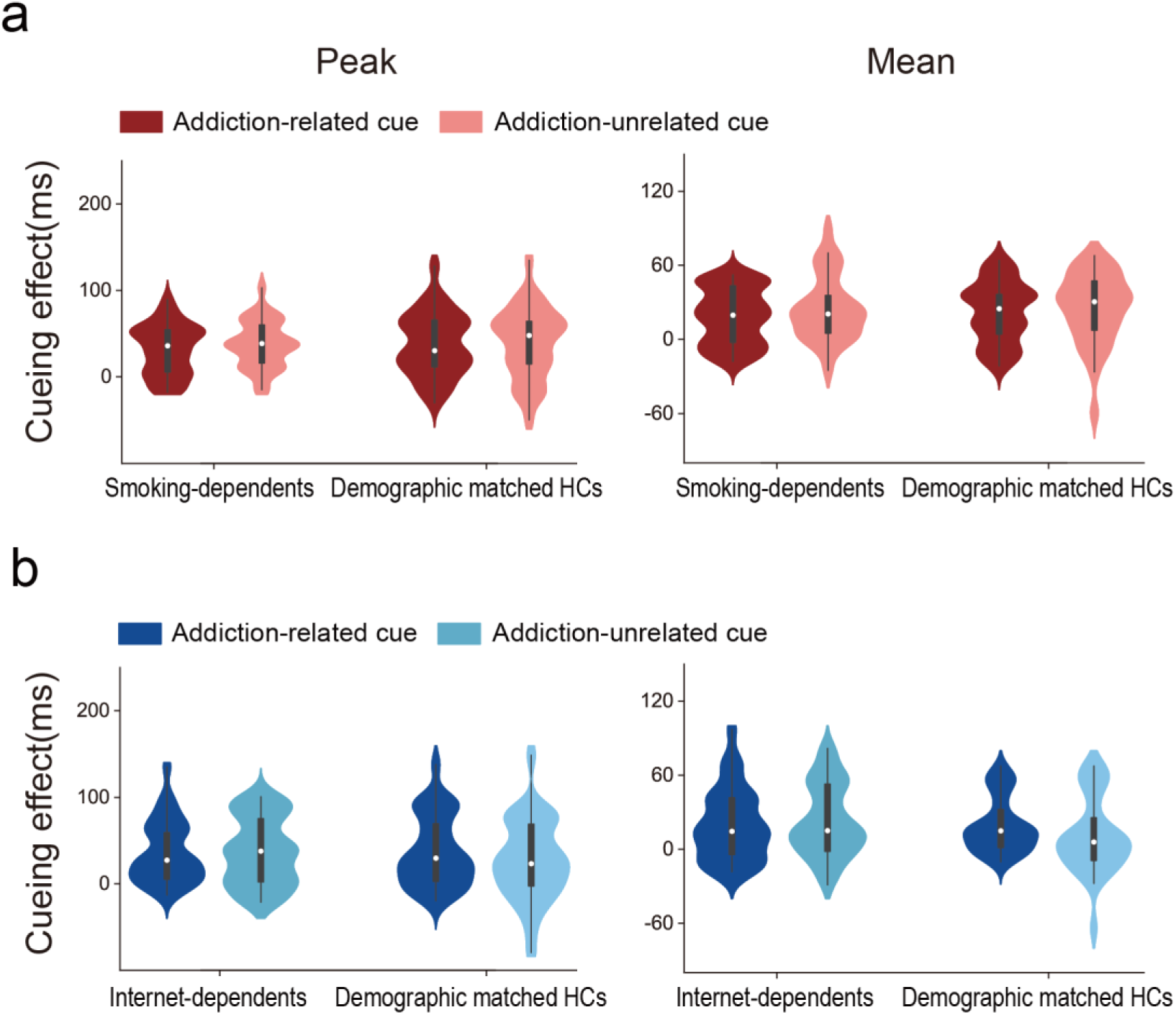
Cueing effects of the test phase. The peak of cueing effect (left), and the mean of cueing effects across distance (right) during addiction-related cue and addiction-unrelated cue conditions, for smoking-dependents (**a**) and internet-dependents (**b**), and their respective HCs. Smoothed density plot denotes the distribution of the sample, box plot denotes the quantile and median scores, and error bars denote 1 SEM calculated across subjects.

### Attention profile

For each group and their HCs, a repeated measures ANOVA with cue (ARC and AUC) and distance (D0-D4) as within-subjects factors showed that, the main effect of cue (Smoking-dependents: F(1, 29) = 1.772, *p* = 0.194, η_p_^2^ = 0.058; HCs of smoking-dependents: F(1, 29) = 0.693, *p* = 0.412, η_p_^2^ = 0.023; Internet-dependents: F(1, 29) = 0.335, *p* = 0.567, η_p_^2^ = 0.011; HCs of internet-dependents: F(1, 29) = 3.833, *p* = 0.060, η_p_^2^ = 0.117) and the interaction between the cue and distance (Smoking-dependents: F(4, 116) = 0.562, *p* = 0.691, η_p_^2^ = 0.019; HCs of smoking-dependents: F(4, 116) = 0.870, *p* = 0.484, η_p_^2^ = 0.029; Internet-dependents: F(4, 116) = 1.019, *p* = 0.400, η_p_^2^ = 0.034; HCs of internet-dependents: F(4, 116) = 0.568, *p* = 0.686, η_p_^2^ = 0.019) were not significant, but the main effect of distance (Smoking-dependents: F(4, 116) = 12.986, *p* < 0.001, η_p_^2^ = 0.309; HCs of smoking-dependents: F(4, 116) = 8.425, *p* < 0.001, η_p_^2^ = 0.225; Internet-dependents: F(4, 116) = 19.928, *p* < 0.001, η_p_ ^2^ = 0.407; HCs of internet-dependents: F(4, 116) = 13.437, *p* < 0.001, η_p_^2^ = 0.317) was significant. Subsequent post hoc paired *t* tests showed that the cueing effect decreased gradually with the distance in smoking-dependents (ARC, D0 vs D1: t(29) = 3.131, *p* = 0.004, Cohen’s d = 0.572; D1 vs D2: t(29) = 0.141, *p* = 0.889, Cohen’s d = 0.026; D2 vs D3: t(29) = 1.136, *p* = 0.041, Cohen’s d = 0.208; D3 vs D4: t(29) =2.144, *p* = 0.041, Cohen’s d = 0.391; AUC, D0 vs D1: t(29) = 3.017, *p* = 0.005, Cohen’s d = 0.551; D1 vs D2: t(29) = 0.573, *p* = 0.571, Cohen’s d =0.105; D2 vs D3: t(29) = 1.546, *p* = 0.133, Cohen’s d = 0.282; D3 vs D4: t(29) = 0.071, *p* = 0.944, Cohen’s d = 0.013, HCs of smoking-dependents (ARC, D0 vs D1: t(29) = 3.086, *p* = 0.004, Cohen’s d = 0.563; D1 vs D2: t(29) = 0.215, *p* = 0.831, Cohen’s d = 0.040; D2 vs D3: t(29) = 1.421, *p* = 0.166, Cohen’s d = 0.259; D3 vs D4: t(29) = 1.375, *p* = 0.180, Cohen’s d = 0.251; AUC, D0 vs D1: t(29) = 1.897, *p* = 0.068, Cohen’s d = 0.347; D1 vs D2: t(29) = 1.495, *p* = 0.146, Cohen’s d = 0.273; D2 vs D3: t(29) = 0.954, *p* = 0.348, Cohen’s d = 0.174; D3 vs D4: t(29) = −0.185, *p* = 0.854, Cohen’s d = −0.034), internet-dependents (ARC, D0 vs D1: t(29) = 5.873, *p* < 0.001, Cohen’s *d* = 1.071; D1 vs D2: t(29) = 1.231 *p* = 0.228, Cohen’s d = 0.225; D2 vs D3: t(29) = 0.019, *p* = 0.985, Cohen’s d = 0.003; D3 vs D4: t(29) = 2.314, *p* = 0.028, Cohen’s d = 0.421; AUC, D0 vs D1: t(29) = 4.296, *p* < 0.001, Cohen’s d = 0.782; D1 vs D2: t(29) = 0.501, *p* = 0.620, Cohen’s d = 0.092; D2 vs D3: t(29) = 1.432, *p* = 0.163, Cohen’s d = 0.261; D3 vs D4: t(29) = 1.589, *p* = 0.123, Cohen’s d = 0.290), and HCs of internet-dependents (ARC, D0 vs D1: t(29) = 2.999, *p* = 0.006, Cohen’s d = 0.548; D1 vs D2: t(29) = 2.878, *p* = 0.007, Cohen’s d = 0.526; D2 vs D3: t(29) = −0.434, *p* = 0.667, Cohen’s d = −0.079; D3 vs D4: t(29) = 1.802, *p* = 0.082, Cohen’s d = 0.329; AUC, D0 vs D1: t(29) = 3.187, *p* = 0.003, Cohen’s d = 0.581; D1 vs D2: t(29) = 2.250, *p* = 0.032, Cohen’s d = 0.410; D2 vs D3: t(29) = 1.454, *p* = 0.157, Cohen’s d = 0.265; D3 vs D4: t(29) = 1.442, *p* = 0.160, Cohen’s d = 0.263) (Fig. 5). These results indicated that for each group, the cueing effect induced by both ARC and AUC conditions was a monotonic gradient profile with a center maximum falling off gradually in the surround.

**Fig. 5.**
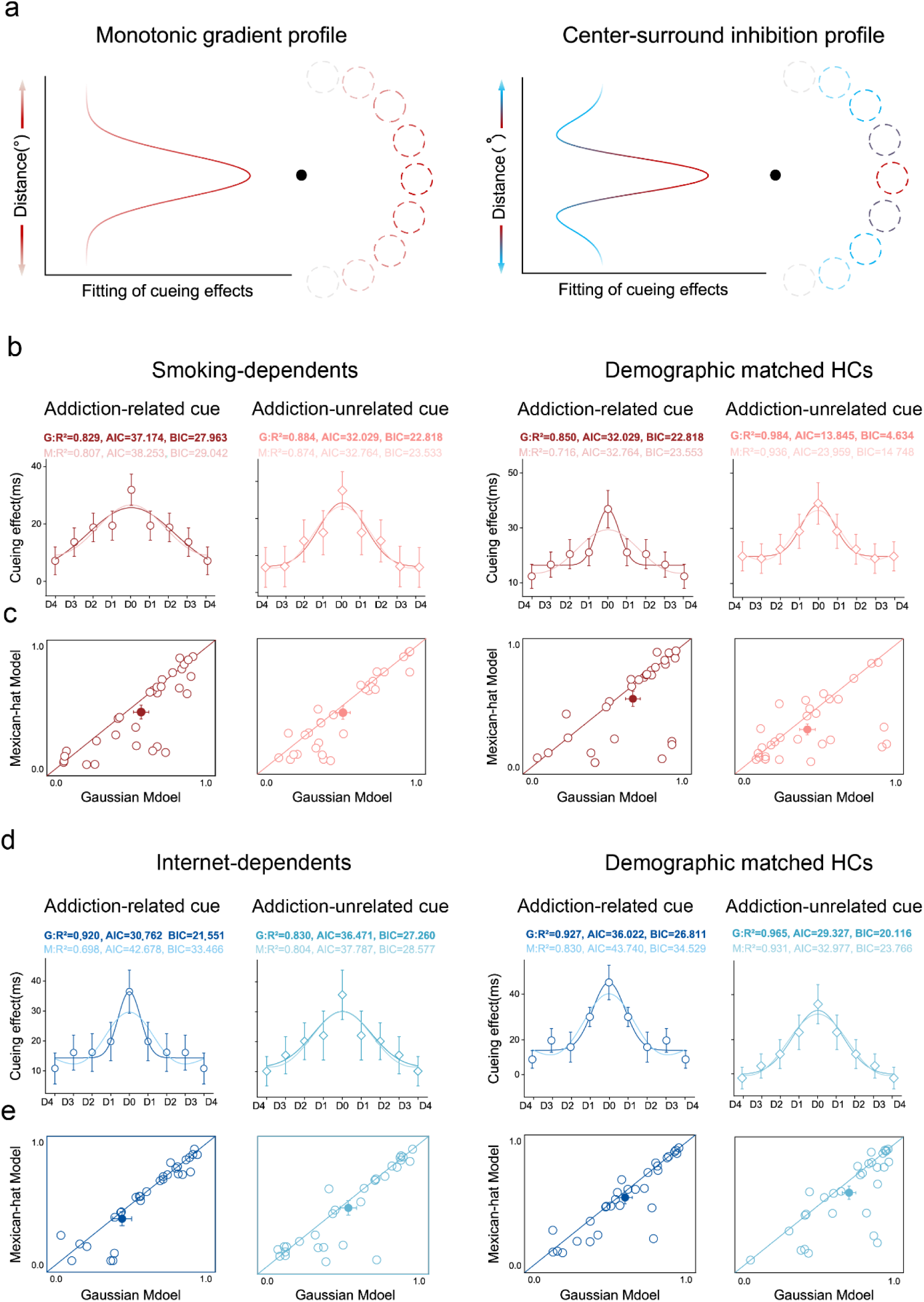
Model fitting and comparison of the cueing effect. a, The potential profile of the cueing effect. Left: the profile of cueing effect displays as a monotonic gradient, with a center maximum falling off gradually in the surround (Gaussian-like). Right: the profile of cueing effect displays a center-surround inhibition, with an inhibitory zone surrounding the focus of bottom-up attention. **b**, The cueing effect of each distance (D0-D4) during addiction-related cue and addiction-unrelated cue conditions, for smoking-dependents (left) and their HCs (right). **c**, *R^2^* of the best fitting Gaussian and Mexican-hat functions of individual subjects for each group and each condition. Most of the dots are below the diagonal line, demonstrating that the Gaussian model was favored over the Mexican-hat model. **d** and **e**, Results of internet-dependents and their HCs, see caption for (**b** and **c**) for a description of each type of graph. Error bars denote 1 SEM calculated across subjects.

To further assess the shape of bottom-up attentional effect, for each group and each condition, we fitted a monotonic model and a nonmonotonic model to the average cueing effect across distances (D0 to D4). The monotonic and nonmonotonic models were implemented as the Gaussian and Mexican-hat (a negative second derivative of a Gaussian function) functions, respectively [7, 51, 71]. To compare these two models to our data, we first computed the Akaike information criterion (AIC) [72] and Bayesian information criterion (BIC) [73] with the assumption of a normal error distribution. Then, we calculated the likelihood ratio (LR) and Bayes factor (BF) of the Gaussian model over Mexican-hat model based on AIC [74] and BIC [75] approximation, respectively. Results showed that, in all groups and all conditions, the LR/BF (see Table 2) strongly favored the Gaussian model over the Mexican-hat model (Fig. 5). Notably, we also conducted similar model comparisons for each subject’ data and found that the Gaussian model was favored over the Mexican-hat model in 18 and 24 out of 30 smoking-dependents (Fig. 5c, left), in 20 and 21 out of 30 HCs for smoking-dependents (Fig. 5c, right), in 21 and 21 out of 30 internet-dependents (Fig. 5e, left), and in 20 and 19 out of 30 HCs for internet-dependents (Fig. 5e, right), during the addiction-related cue (ARC) and addiction-unrelated cue (AUC) conditions, respectively. In addition, across individual subjects, a non-parametric Wilcoxon signed-rank test was conducted to compare the *R^2^* of two models, and results significantly advocated the Gaussian model over the Mexican-hat model on each group and each condition (Smoking-dependents, ARC: z = 2.306, *p* = 0.011, effect size: r = 0.421; AUC: z = 2.629, *p* = 0.005, r = 0.480; HCs of smoking-dependents, ARC: z = 1.656, *p* = 0.049, r = 0.302; AUC: z = 2.078, *p* = 0.019, r = 0.379; Internet-dependents, ARC: z = 1.581, *p* = 0.057, r = 0.289; AUC: z = 1.779, *p* = 0.038, r = 0.325; HCs of internet-dependents, ARC: z = 2.087, *p* = 0.018, r = 0.381; AUC: z = 2.303, *p* = 0.011, r = 0.420). Together, these results further constituted strong evidence for the monotonic gradient profile of bottom-up attention with and without addiction, suggesting that the attention profile also cannot demarcate either internet-dependents or smoking-dependents from arbitrary rewards.

**Table 2.** LR/BF of the model comparison for each group and each condition.

| Gaussian model over Mexican-hat model | ARC | AUC |
| --- | --- | --- |
| Smoking-dependents | $1.72 \times 10^0$ | $1.44 \times 10^0$ |
| HCs of smoking-dependents | $1.74 \times 10^1$ | $1.57 \times 10^2$ |
| Internet-dependents | $3.87 \times 10^2$ | $1.93 \times 10^0$ |
| HCs of internet-dependents | $4.74 \times 10^1$ | $6.20 \times 10^0$ |

### Attention field

Our results indicated that, for each group and each condition, the spatial focus of bottom-up attention was best explained by the Gaussian rather than Mexican-hat model. To quantitatively examine the scope of bottom-up attentional modulation, we fitted the cueing effects from D0 to D4 with a Gaussian function and used the FWHM (full width at half maximum) bandwidth of the Gaussian to quantify their scopes (attention fields). For each addiction group and their HCs, the fitted FWHM bandwidths were submitted to a two-factor mixed ANOVA with group (dependents and HCs) as the between-subjects factor and cue (ARC and AUC) as the within-subjects factor. Results showed that, for smoking-dependents and their HCs, the main effect of group (F(1, 58) = 0.210, *p* = 0.648, η_p_^2^ = 0.004), the main effect of cue (F(1, 58) = 2.228, *p* = 0.141, η_p_^2^ = 0.037), and the interaction between these two factors (F(1, 58) = 0.003, *p* = 0.953, η_p_^2^ < 0.001) were not significant (Fig. 6a). For internet-dependents and their HCs, the main effect of group was not significant (F(1, 58) = 0.400, *P* = 0.530, η_p_^2^ = 0.007), but the main effect of cue (F(1, 58) = 8.569, *p* = 0.005, η_p_^2^ = 0.129) and the interaction between these two factors (F(1, 58) = 5.718, *p* = 0.020, η_p_^2^ = 0.090) were both significant. Further simple effect analysis showed that the FWHM bandwidth of ARC condition was significantly smaller than that of AUC condition for internet-dependents (t(29) = −3.867, *p* = 0.001, Cohen’s d = −0.706), but not for their HCs (t(29) = −0.369 *p* = 0.715, Cohen’s d = −0.067); the FWHM bandwidth of internet-dependents was significantly smaller than that of HCs for ARC condition (t(58) = −2.227, *p* = 0.027, Cohen’s d = −0.706), but not for AUC condition (t(58) = 0.771, *p* = 0.444, Cohen’s d = 0.202) (Fig. 6b). These results indicate a reduction in the AF of bottom-up attention in internet- but not smoking-dependents, and therefore, the AF could as a cognitive-behavioral marker for demarcating internet-addiction from arbitrary rewards.

**Fig. 6.**
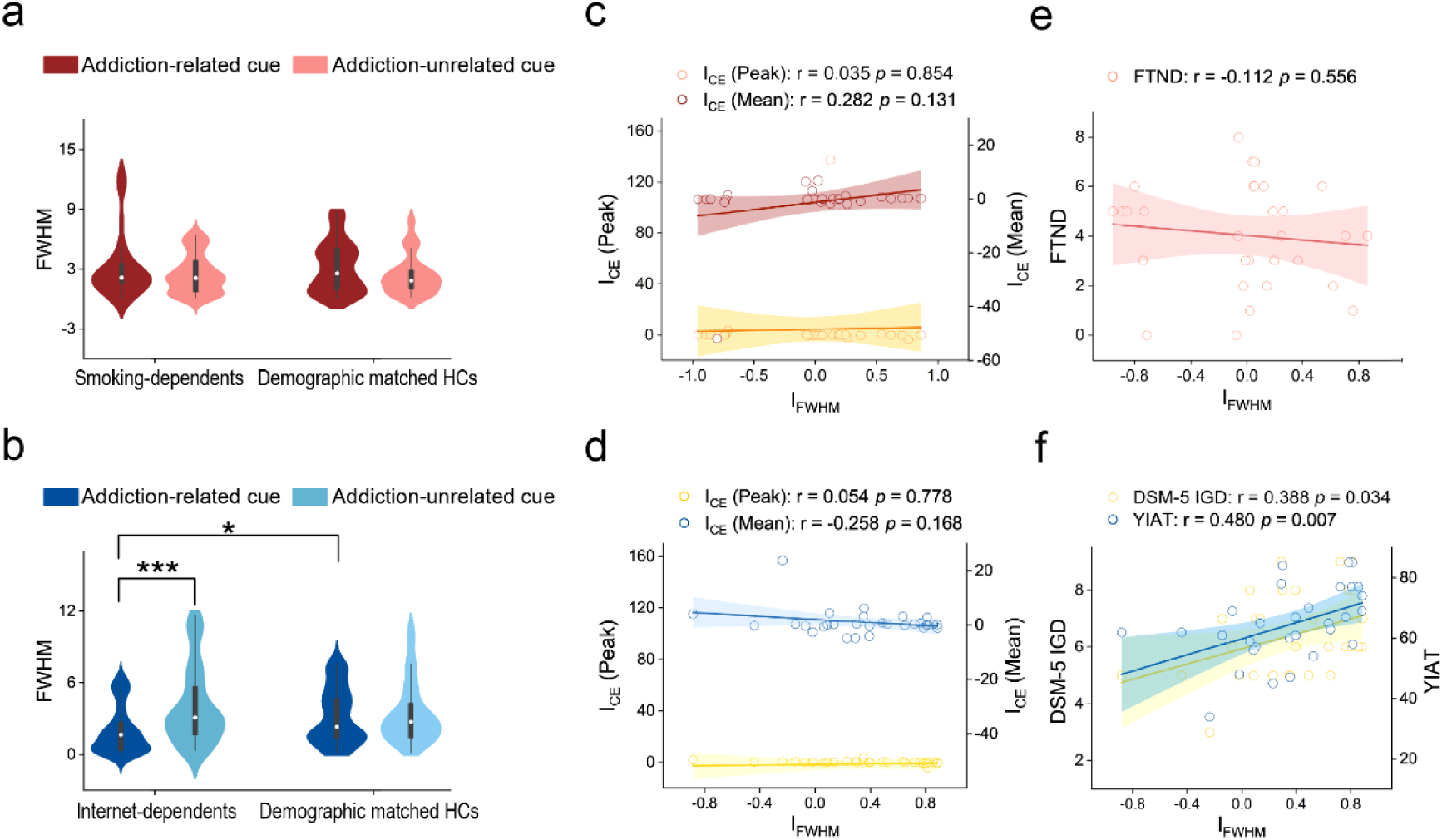
Fitted FWHM bandwidth of the Gaussian function for each group and each condition, and results of correlation analysis. The fitted FWHM bandwidth of monotonic Gaussian model during addiction-related cue (ARC) and addiction-unrelated cue (AUC) conditions for smoking-dependents (**a**), internet-dependents (**b**), and their respective HCs. Smoothed density plot denotes the distribution of the sample, box plot denotes the quantile and median scores, and error bars denote 1 SEM calculated across subjects. Correlations between the decreased FWHM bandwidth and peak cueing effect, and between the decreased FWHM bandwidth and mean cueing effect across distances, in ARC relative to AUC, across individual smoking-dependents (**c**) and internet-dependents (**d**). **e**, Correlations between the decreased FWHM bandwidth and FTND across individual smoking-dependents. **f**, Correlations between the decreased FWHM bandwidth and DSM-5 IGD, and between the decreased FWHM bandwidth and YIAT, across individual internet-dependents.

Notably, one could argue that within each group, our observed AF results (either significant or non-significant) between ARC and AUC conditions could be explained by their differences in cueing effect. To examine this issue, for each group, we first computed an AF index I_FWHM_ to quantify how much the fitted FWHM bandwidth decreased during the ARC condition relative to the AUC condition. The index was calculated as follows: I_FWHM_ = (FWHM_AUC_ - FWHM_ARC_) / (FWHM_AUC_ + FWHM_ARC_) * 100%, where FWHM_AUC_ and FWHM_ARC_ are the fitted FWHM bandwidth in the ARC and AUC conditions, respectively. Similarly, we also computed a cueing effect index I_CE_ to quantify how much the cueing effect decreased during the ARC condition relative to the AUC condition. The index was calculated as follows: I_CE_ = (CE_AUC_ - CE_ARC_) / (CE_AUC_ + CE_ARC_) * 100%, where CE_AUC_ and CE_ARC_ are the measured cueing effect (either the peak or the mean of cueing effects) in the ARC and AUC conditions, respectively. Then, we calculated the correlation coefficients between the I_FWHM_ and I_CE_ across individual subjects. If the comparison of AF between ARC and AUC condition is derived by their differences in cueing effect, then we would observe a significant correlation between these two indexes across individual subjects. For each group, however, the I_FWHM_ was not significantly correlated with the I_CE_ for either the peak of cueing effect (Smoking-dependents: r = 0.035, *p* = 0.854; HCs of smoking-dependents: r = -0.335, *p* = 0.070; Internet-dependents: r = 0.054, *p* = 0.778; HCs of internet-dependents: r = 0.213, *p* = 0.259) or the mean of cueing effects across distances (Smoking-dependents: r = 0.282, *p* = 0.131; HCs of smoking-dependents: r = −0.239, *p* = 0.203; Internet-dependents: r = −0.258, *p* = 0.168; HCs of internet-dependents: r = 0.243, *p* = 0.195) (Fig. 6c and 6d), which strongly against the cueing effect explanation.

Importantly, to examine whether the reduction in AF closely mirrored addictive symptom severities, across individual subjects, we calculated the correlation coefficients between I_FWHM_ and FTND for smoking-dependents, and between I_FWHM_ and DSM-V IGD/ YIAT for internet-dependents. Results showed that the I_FWHM_ correlated significantly with both the DSM-V IGD (r = 0.388, *p* = 0.034) and YIAT (r = 0.481, *p* = 0.007) across internet-dependents (Fig. 6f), but not with FTND across smoking-dependents (r = −0.112, *p* = 0.556, Fig. 6e). These results further confirm a unique AF alteration in internet- but not smoking-dependents, and the AF could be considered as a cognitive-behavioral marker for demarcating internet-addiction from arbitrary rewards.

Across different groups, to further against the cueing effect explanation, we directly compared the FWHM bandwidth between the addiction group and their HCs during both ARC and AUC conditions, with the cueing effect (either the peak or mean) as the covariate. Results confirmed that, for smoking-dependents, their FWHM bandwidths did not differ from HCs in either ARC (the peak of cueing effect as the covariate: F(1, 57) = 0.060, *p* = 0.807, η_p_^2^ = 0.001 ; the mean of cueing effects as the covariate: F(1, 57) = 0.126, *p* = 0.724, η_p_^2^ = 0.002) or AUC (the peak of cueing effect as the covariate: F (1, 57) = 0.177, p = 0.676, η_p_^2^ = 0.003; the mean of cueing effects as the covariate: F (1, 57) = 0.155, p = 0.695, η_p_^2^ = 0.003) conditions. However, the FWHM bandwidth of internet-dependents was significantly smaller than that of HCs for ARC condition (the peak of cueing effect as the covariate: F (1, 57) = 5.215, *p* = 0.026, η_p_^2^ d = 0.084; the mean of cueing effects as the covariate: F (1, 57) = 5.101, *p* = 0.028, η_p_^2^ = 0.082), but not for AUC condition (the peak of cueing effect as the covariate: F (1, 57) = 0.487, *p* = 0.488, η_p_^2^ = 0.008; the mean of cueing effects as the covariate: F (1, 57) = 0.170, *p* = 0.682, η_p_^2^ = 0.003). Altogether, these results further confirm a unique AF alteration in internet- but not smoking-dependents, and this finding was not explained by the difference in cueing effect.

### Divisive-normalization model simulations

Our results can be viewed as identifying a narrowed scope of addiction-related attentional bias in internet-dependents, in other words, a sharpened gradient of addiction-related attentional bias, also known as the “tunnel vision”. Parallel to the classical contrast response function [7, 10, 76] in divisive-normalization model of attention, the population gain can be considered as a function of probe distance (D0-D4) from the exogenous cue. For internet-dependents, the gradient of the population gain should be steeper for ARC than AUC conditions. Therefore, we simulated our empirical data with the divisive-normalization model of attention using custom MATLAB scripts based on the code of Reynolds and Heeger [48] with four free parameters: the gain of attention [*A(x, u)*], separately optimized for each group and each condition, the normalization constant *σ*, the orientation tuning of AF, and a scaling parameter to linearly scale simulated values to performance (the cueing effect) (Fig. 7a). Results showed that for internet-dependents, the simulated AF in the ARC condition was significantly smaller than that in the AUC condition (t(29) = -2.598, *p* = 0.015, Cohen’s d = - 0.965, Fig. 7b). Similarly, we computed a simulated AF index I_AF_ to quantify how much the fitted AF decreased during the ARC condition relative to the AUC condition. The index was calculated as follows: I_AF_ = (AF_AUC_ - AF_ARC_) / (AF_AUC_ + AF_ARC_) * 100%, where AF_AUC_ and AF_ARC_ are the fitted AF in the ARC and AUC conditions, respectively. Then, we calculated the correlation coefficients between I_AF_ and DSM-V IGD/ YIAT for internet-dependents. Results showed that across internet-dependents, the I_AF_ correlated significantly with the YIAT (r = 0.516, *p* = 0.004), but not with the DSM-V IGD (r = 0.173, *p* = 0.360) (Fig. 7c), further confirming a unique AF alteration in internet-dependents.

**Fig. 7.**
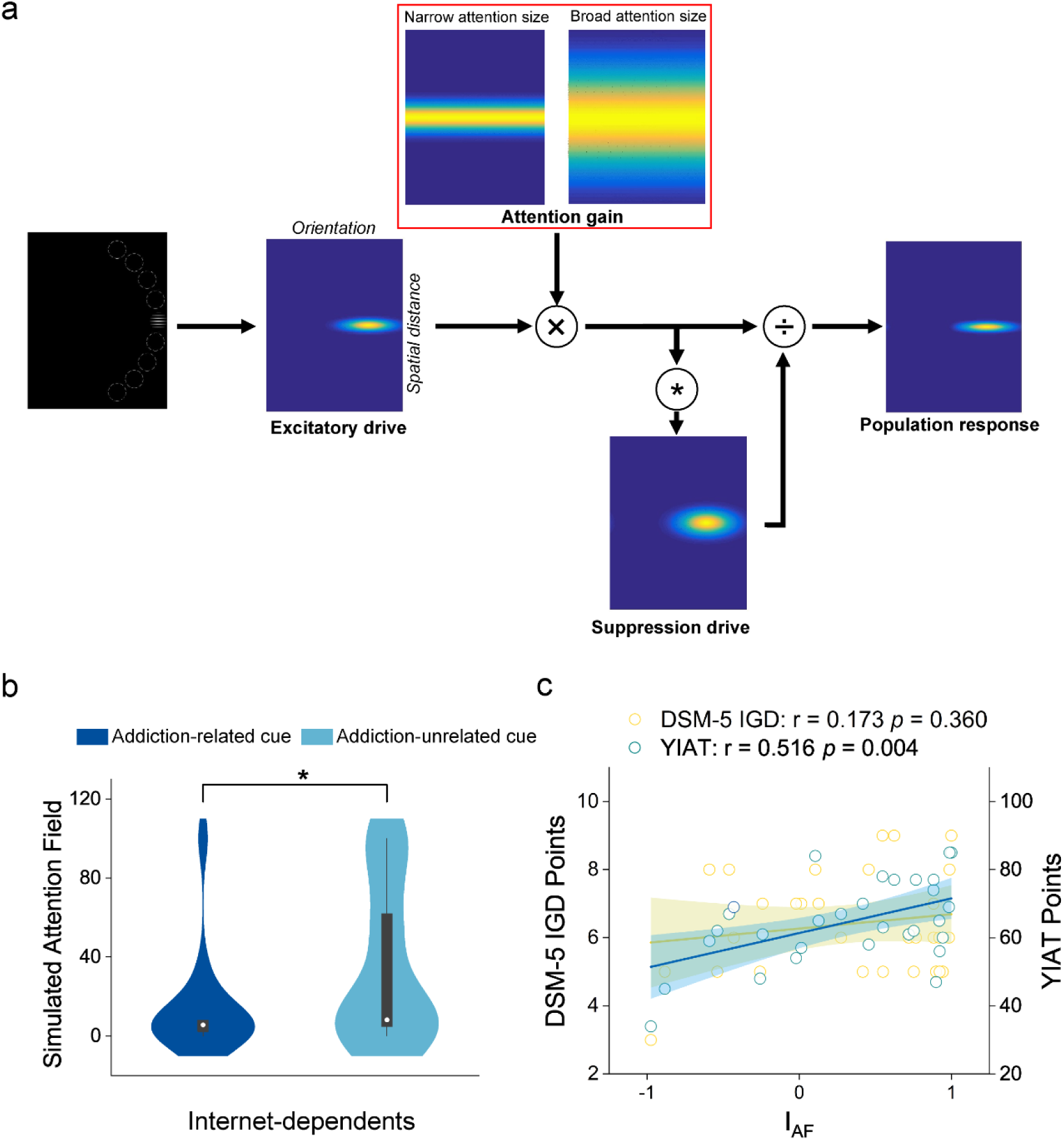
Simulated attention fields in the Divisive-normalization model of attention. a, The normalization model diagram [48]: narrow vs. broad attention size was modeled using narrower vs. broader spatial Gaussians in the gain of attention [*A(x, u)*], optimized for addiction-related and addiction-unrelated cue conditions, respectively. **b,** Simulated attention field *A(x, u)* between addiction-related and addiction-unrelated cue conditions. Smoothed density plot denotes the distribution of the sample, box plot denotes the quantile and median scores, and error bars denote 1 SEM calculated across subjects. **c,** Correlations between the decreased *A(x, u)* and two questionnaires, cross individual internet-dependents.

### Covariate analyses with demographic characteristics

To exclude potential fluences of demographic characteristics on our results, using the same two-factor mixed ANOVA with group (addiction group and HCs) as the between-subjects factor and cue (ARC and AUC) as the within-subjects factor, we re-analyzed both the AE and FWHM bandwidth with each of the demographic characteristics as the covariate. The covariate could be age, depression, IQ, DSM-5 IGD, or YIAT for smoking-dependents and their HCs, and age, depression, or IQ for internet-dependents and their HCs. Results showed that all demographic covariates did not appear to have an impact on either the AE or FWHM bandwidth results (see Table S1), further confirming our conclusion that the AF could be considered as a cognitive-behavioral marker for demarcating internet- but not smoking-addiction from arbitrary rewards.

## Discussion

We probed addiction-related bottom-up attentional bias, including its attentional effect (AE), attention profile (AP), and attention field (AF), free from reward-driven attentional captures and found the following psychophysical results. First, for both internet-dependents and smoking-dependents, we found support for previous studies arguing that the AE of addiction could reflect a normal cognitive process that promotes reward-seeking behavior [27, 29, 39, 41, 77]. This addiction-related AE distinguishes neither between individuals with and without a history of addiction, nor between addiction-related and addiction-unrelated stimuli when previously associated with arbitrary rewards, such as monetary in our study (Fig. 4). Second, we further found that the AP of each group and each condition displayed as a monotonic gradient profile, with a center maximum falling off gradually in the surround (Gaussian-like, Fig. 5), indicating that the AP also cannot demarcate either internet-dependents or smoking-dependents from arbitrary rewards. Third, however, we found a unique AF alteration in internet- but not smoking-dependents (Fig. 6), and intriguingly, the decreased AFs in internet-dependents can be simulated by using the classical divisive-normalization model of attention [48]. More importantly, across internet-dependents, both the observed and simulated reductions of AFs closely tracked their addictive symptoms severities (Fig. 7), suggesting that the AF can be considered as a cognitive-behavioral marker for demarcating internet-addiction from arbitrary rewards. Finally, our results cannot be explained by the eye movement (Fig. S3), the degree of association between stimuli and rewards (Fig. 2), or demographic variables (Table S1), as in all these factors, there were no significant differences/influences among conditions.

### Gradient profile of bottom-up attentional bias with and without addiction

In accordance with previous neurophysiological [78–80], psychophysical [7, 52, 81–86], eye movement [53], electroencephalographic [52, 87, 88], and functional magnetic resonance imaging [54] studies, our study found that for each group and each condition, the bottom-up attention-triggered bias was a monotonic gradient profile with a center maximum falling off gradually in the surround (Gaussian-like, Fig. 5). However, a number of neurophysiological [80, 89, 90], psychophysical [91–93], and brain imaging [49,50,94–96] studies, as well as a computational model [97, 98] have conversely reported a center surround (Mexican hat-like) profile where a zone of sensory attenuation surrounds a center region of facilitation. We suggested that this striking discrepancy might be due to two different factors. One is the experimental task or paradigm. Specifically, for example, in Hopf et al. [49] where inhibition was seen surrounding the locus of spatial attention, subjects were asked to search for a target (i.e., the exogenous cue in our study), which appeared randomly to change the spatial focus of attention and thus was task-relevant. While they measured the event-related magnetic field response elicited by a task-irrelevant probe that appeared or was absent with equal probability. In our study, by contrast, we used a modified Posner cueing paradigm to measure the probe’s attentional effect induced by the exogenous cue, either addiction-related or addiction-unrelated. Subjects were asked to discriminate the orientation (45° or 135°) of the probe (Fig. 3c); thus, the exogenous cue was task-irrelevant while the probe was task-relevant. Directing attention to the target probe in our study here could conflate perceptual and post-perceptual mechanisms of attention, thus eliminating or strongly attenuating this suppression effect.

The other that may influence the spatial profile of bottom-up attention is whether the exogenous cue was presented with (e.g., Hopf et al. [49]) or without (e.g., the current study) the distractors. As is known to all, the biased competition model of attention [99] proposes that attention operates when multiple stimuli compete for access to neural representation, and this competition occurs when multiple stimuli fall within a neuron’s receptive field. In this case, distractors within a receptive field are suppressed while attended stimuli are enhanced. Additionally, Tsotsos and colleagues [97, 98] proposed the selective tuning model, which directly suggested that the inhibitory zone surrounding the attended item results from top-down propagation of a winner-take-all mechanism that attenuates irrelevant upstream connections iteratively from one hierarchical level down to the next. Thus, without the distractor, these irrelevant upstream connections across the visual cortical processing hierarchy could be eliminated or strongly attenuated, which, in turn, affects the inhibitory zone surrounding the attended target, in other words, resulting in a monotonic gradient rather than center-surround profile of attentional modulation. Therefore, further work is needed using different paradigms (e.g., with and without distractors) to confirm the spatial profile of bottom-up attentional bias with and without addiction.

### A narrowed scope of bottom-up attentional bias in internet- but not smoking-dependents

Our findings can be viewed as identifying an addiction-dependent scope of bottom-up attentional bias for internet-dependents. Note that this conclusion is based on a report-based paradigm in which subjects overtly push a button to report their percept. Several studies have argued that such report-based paradigms could be modulated by factors that are not directly related to the scope of attentional modulation, such as higher-level strategies, response history, experience, learning, response biases, and personality [100]. In addition, using a no-report paradigm, such as recording subjects’ pupillary light responses, Tkacz-Domb and Yeshurun [53] revealed that the scope of attentional modulation was twofold larger than that estimated using the traditional report-based paradigm. It is important to note that subjects in our study performed exactly the same task between addiction-related and addiction-unrelated cue conditions, and between internet-dependents and HCs (Fig. 3c), thus the addiction-dependent scope of bottom-up attention evident here cannot be explained by this discrepancy.

Intriguingly, previous studies have reported different AFs relative to healthy controls in several psychiatric conditions, such as autism spectrum disorders [101–103], anxiety [104], and depression [105]. Here, our results extend the cognitive-behavioral marker of AFs into the internet-addiction, or, more generally, non-substance addictions.

A narrowed scope of bottom-up attention in internet-addictions not only supported previous studies proposing that the narrowing of the attention field is sometimes referred to as “weapon focus”, in which peripheral details of stimuli are more poorly encoded, as measured in later memory [106], repeated adaptation [107], emotional attention [10], and unconscious attention [7], but also confirmed several studies reporting that the internet-addiction could enhance both the spatial resolution [108] and selection [109] of attention. More importantly, our study succeeded in linking this narrowed scope of bottom-up attention in internet-addictions with the classical divisive-normalization model. Although the physiological roots of this link have never been known, our results could infer an imbalance between the spatial extent of both low-level (receptive field) and higher-order (attentional signal) components of the addicted visual system [48]. People with internet-addictions here could be described as having tunnel vision, attracted to details of a visual scene while neglecting surrounding stimuli, as if attention were sharply pinpointed to the peaks of their visual world. This finding not only provides insight into mechanisms underlying how individuals with internet-addiction perceive the world, but also a starting point in a well characterized domain for modeling atypical neural circuitry in the addicted brain.

### Non-substance addiction is not an ill-posed addiction

Some theories of addiction propose that each addiction – whether it be to substance- or non-substance-related – might be a distinctive expression of the same underlying syndrome. In other words, addiction is a syndrome with multiple opportunistic expressions [110, 111]. Indeed, previous studies have demonstrated that both substance- and non-substance-related addictions share common genetic polymorphisms [112, 113], predisposing factors [114], pathological characteristics [115], associated ramifications [116], neuronal and biochemical underpinnings [114, 117, 118]. However, several studies argued that non-substance-related addictions were quite different from substance-related addictions in clinical manifestation [119], more akin to an impulse-control disorder than substance-related addictions [120], often co-occurred with other psychiatric disorders and would be cured naturally after treatments of these disorders [121]. Non-substance-related addictions may simply be an inappropriate-coping [122] or excessive [123] behavior, without significant impairment of social functions [124], that doesn’t fall under the medical concept of ‘addiction’ [125]. Conversely, some of non-substance-related addictions, such as the internet gaming disorder could enhance subjects’ contrast sensitivity [126], balance rehabilitation [127], memory/working memory [128, 129], spatial cognition [108, 130], visual selective attention [109], the accessibility of prosocial thought [131], and various cognitive performance [132]. Therefore, some of researchers have argued that non-substance-related addictions could be not real addictions [133]. Intriguingly, however, our study identifies the AF as a cognitive-behavioral marker for demarcating internet-dependents, one of non-substance-related addictions, but not smoking-dependents from arbitrary rewards, arguing against this ill-posed notion.

Although our study failed to demarcate smoking-dependents from arbitrary rewards, we cannot generalize this conclusion to other substance-related addictions, such as drugs or alcohols. Several studies have found that smoking-dependents, when compared to other substances, was associated with equal or greater levels of difficulty in quitting and urge to us [134], but that it was not as pleasurable [135] and did not cause behavioral harms [136]. Conversely, most smoking-dependents describe their subjective effects of smoking in terms of improving their functioning, i.e. improved concentration and decreased stress, anxiety, depression, or anger [137]. In addition, drug-dependents produce an overflow of dopamine outside the nucleus accumbens that is associated with euphoria and potentiates drug reinforcement, whereas this effect is very small with smoking-dependents [138]. More importantly, some of the generic dependence criteria do not apply to smoking-dependents and some well-validated measures of smoking-dependents are not included in the generic criteria [139]. Given these dissimilarities between smoking-dependents and other substances, several studies have argued that smoking-dependents might be not really a substance addiction [140, 141]. Further work is thus needed to use various substance-related addictions to address the relationship between addictions and rewards.

### Conclusions

In sum, our study provides, to the best of our knowledge, the first evidence revealing a unique AF alteration in internet- but not smoking-dependents. Identifying the AF as a cognitive-behavioral marker for demarcating internet-addiction, or, more generally, non-substance addiction, from arbitrary rewards, not only gives insight into the potential cross contamination between addiction and reward in controlling the nature of attention selection, but also further argues against the notion that the non-substance addiction is ill-posed.

## Methods

### Subjects

A total of four groups participated in the study: 30 smoking-dependents, 30 demographic-matched HCs of smoking-dependents, 30 internet-dependents, and 30 demographic-matched HCs of internet-dependents. They were recruited through advertisements. All subjects were asked to refrain from their addiction-related stimuli (smoking for smoking-dependents and their HCs; internet for internet-dependents and their HCs) 4–6 h prior to the experiment. All of them participated in the training and test phases. All subjects were naïve to the purpose of the study. They reported normal or corrected-to-normal vision and had no known neurological, psychiatric, or visual disorders. They gave written informed consent in accordance with protocols approved by the human subjects review committee of School of Psychology at South China Normal University. Sample characteristics and addiction-related measures are outlined in Table 1.

### Psychometric Testing

All subjects completed the standard Raven progressive matrices (RPM) [61], Beck Depression Inventory-II (BDI-II) [60], Fagerström Test for Nicotine Dependence (FTND) [57], Internet gaming disorder (IGD) in Section 3 of the DSM-V [62], and the Young’s Internet Addiction Test (YIAT) [63]. BDI-II, whose Cronbach’s alpha was 0.92, is a 21-item self-administered questionnaire assessing the severity of depressive symptoms, with higher scores indicates the severer depressive symptoms. FTND, whose Cronbach’s alpha was 0.89, is a widely used 6-items, self-report measure of nicotine dependence. The possible range of scores is from 0-10, with 5 indicating moderate nicotine dependence. DSM-V IGD, whose Cronbach’s alpha was 0.93, is composed by 9 items on yes or no scale, with 5 indicating gaming disorder. YIAT, whose Cronbach’s alpha was 0.84, is composed by 20 items on five-point Likert scale, with 50 points indicating internet addiction.

### Stimuli

In the training phase, for each dependent group (internet-dependents and smoking-dependents) and their respective demographic-matched HCs, there were 5 addiction-related and 15 addiction-unrelated images. All images were scaled to the same size (diameter: 1.95°). On each trial, a pair of images was randomly centered in the upper and lower hemifields at 6.28° eccentricity, one of which was rewarded images. For the addiction-related reward condition, the rewarded stimuli were 5 addiction-related images. For the addiction-unrelated reward condition, the 5 rewarded stimuli were chosen randomly from 15 addiction-unrelated images. The remaining 10 addiction-unrelated images were randomly and equally divided into two groups, as the unrewarded stimuli for each condition (Fig. 1a). In the test phase, as illustrated in Fig. 3b, each texture stimulus contained 18 positions settled at an iso-eccentric distance from fixation (9.47° of visual angle): half of them were located in the left visual field, and the other half were located in the right visual field. The center-to-center distance between two neighboring positions was 2.50°. For the cue frame, a pair of stimuli always appeared in the center of 9 positions, with a pre-rewarded addiction-related image and an unrewarded image (i.e., the addiction-related cue condition), a pre-rewarded addiction-unrelated image and an unrewarded image (i.e., the addiction-unrelated cue condition), or two unrewarded images (i.e., the non-cue condition). The pre-rewarded stimulus, either addiction-related or addiction-unrelated, served as an exogenous cue to attract subjects’ covert spatial attention (Fig. 3a). The probe grating (spatial frequency: 4.0 cycles/°; diameter: 1.95°; phase: random) was oriented at 45° or 135° away from the vertical and appeared equiprobably and randomly across the possible 9 positions. Thus, there were five possible distances between the exogenous cue and probe, ranging from D0 (cue and probe at the same location) through D4 (cue and probe four items away from each other) (Fig. 3b).

### Training phase Procedure

In the training phase, for each dependent group and their demographic-matched HCs, subjects performed a probabilistic instrumental learning task with monetary outcomes, adapted from previous studies [64–67]. Each of the pairs of stimuli was associated with pairs of outcomes (gain 1 ¥ or 0 ¥), the two stimuli corresponding to reciprocal probabilities (0.8/0.2 and 0.2/0.8). For addiction-related reward conditions, the probabilities of winning 1 ¥ were 0.8/0.2 for the addiction-related image and 0.2/0.8 for the addiction-unrelated image. For addiction-unrelated reward conditions, the probabilities of winning 1 ¥ were 0.8/0.2 for one of the addiction-unrelated images and 0.2/0.8 for the other. For both addiction-related and addiction-unrelated reward conditions, each pair was randomly presented 25 times for a total of 125 trials. On each trial, one pair was randomly presented and the relative position of two images was counterbalanced across trials. Subjects were required to choose either the upper or lower stimulus by pressing “*I*” or “*J*”, respectively. Then, the choice was circled in red and the outcome was displayed on the screen, gain either 1 ¥ or 0 ¥ (Fig. 1b). Therefore, to win money, subjects had to learn, by trial and error, the stimulus–outcome associations. They were told that they would play for real money, but at the end of the experiment their winnings were rounded up to a fixed amount.

### Computational model

We used a standard *Q*-learning algorithm [64, 68–70] to fit the reinforcement learning for each subject’s sequence of choices. For each pair of stimuli and each group of subjects, such as an addiction-related image *a* and an addiction-unrelated image *b*, the model estimates the expected values of choosing *a* (*Qa*) and choosing *b* (*Qb*), based on individual sequences of choices and outcomes. This value, termed a *Q* value, is essentially the expected reward obtained by taking that particular action. These *Q* values were set at 0 before learning, and after every trial (t > 0) the value of the chosen stimulus (for example, the addiction-related image *a*) was updated according to the rule:

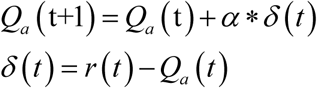

where *^Qa^* ^(t)^ and *^Qa^* ^(t+1)^ is the expected outcome for the *t* and *t+1* trials, respectively; α is the learning rate that fits best to each subject’s choice behavior reflects the degree to which previous reinforcement outcomes affect subsequent *Q* values; *δ* (*t*) is the outcome prediction error, calculated as the difference between the actual *r* (*t*) and the expected *^Qa^* ^(t)^ outcome. In our study, the reinforcement *r* (*t*) was either 1 ¥ or 0 ¥. Given the expected *Q* values, the probability *P_a_* (*t*) of the observed choice was estimated with the softmax rule:

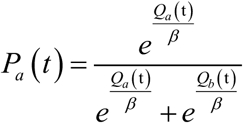

where *β*, the softmax temperature, reflects the stochasticity of the softmax function. Thus, higher *β* values indicate exploration (i.e., choosing the lower *Q*-value option) and lower *β* values indicate exploitation (i.e., choosing the higher value option). The constants *α* (learning rate) and *β* (temperature) were adjusted to maximize the likelihood of the actual choices under the model, across all conditions.

### Test phase

#### Procedure

The test phase consisted of two sessions (the addiction-related and addiction-unrelated), which were the same except for the exogenous cue: addiction-related or addiction-unrelated. Subjects performed two sessions on the same day, and the order of the two sessions was counterbalanced across subjects. Each session consisted of 16 blocks of 80 trials, 40 for the left visual field and 40 for the right visual field. In each block and each visual field, each trial began with the fixation. A cue frame with (the cue condition) or without (the non-cue condition) exogenous cue was equiprobably and randomly presented for 50 ms, followed by a 150 ms fixation interval. Then a probe grating, orientating at 45° or 135° away from the vertical, was presented for 50 ms. Subjects were asked to press one of two buttons as rapidly and correctly as possible to indicate the orientation of the probe (45° or 135°) (Fig. 3c). The cueing effect for each distance (D0-D4) was quantified as the difference between the reaction time of the probe task performance in the non-cue condition and that in the cue condition. One should note that our study cannot design the valid and invalid cue conditions developed by the classical Posner cueing paradigm to measure the cueing effect of each distance (D0-D4) since the distance (visual angle) between the exogenous cue and probe was varied not only in the valid cue condition but also in the invalid cue condition^7^.

#### Model fitting and comparison

For each subject and each condition, we fitted a monotonic model and a non-monotonic model to averaged cueing effect. The monotonic and non-monotonic models were implemented as the Gaussian and Mexican-hat (i.e., a negative second derivative of a Gaussian function) functions [7, 51, 71], respectively, as follows:

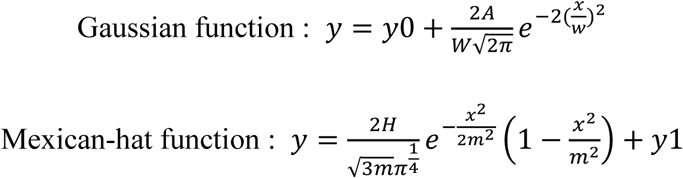

where *y* is the cueing effect, *x* is the distance; *w*, *A*, and *y0* are the three parameters controlling the shape of the Gaussian function; *m*, *H*, and *y1* are three parameters controlling the shape of the Mexican-hat function. To compare these two models to our data, we first computed the Akaike information criterion (*AIC*) [72] and Bayesian information criterion (*BIC*) [73], with the assumption of a normal error distribution as follow:

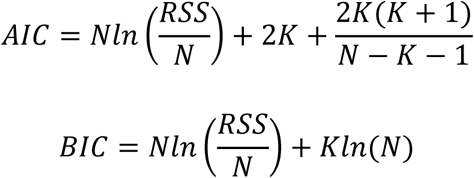

where *N* is the number of observations, *K* is the number of free parameters, and *RSS* is residual sum of squares. Then, we further calculated the likelihood ratio (*LR*) and Bayes factor (*BF*) of the Gaussian model over the Mexican-hat model based on *AIC* [74] and *BIC* [75] approximation, respectively, as follows;

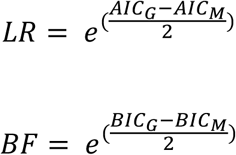

where *AIC_G_* and *BIC_G_* are for the Gaussian model and *AIC_M_* and *BIC_M_* are for Mexican-hat models. The results indicated that, for each group and each condition, the Gaussian model was strongly favored over the Mexican-hat model (Fig. 5). Thus, to quantitatively examine the AF of attentional modulation, we fit the averaged cueing effects from D0 to D4 with a Gaussian function and used the FWHM (full width at half maximum) bandwidth of the Gaussian to quantify their attention fields:

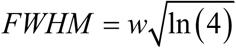

where *w* is the fitted width of the Gaussian function.

#### Divisive-normalization model simulations

The normalization model of attention [48] computes the response of an arbitrary single neuron to a given set of stimuli as:

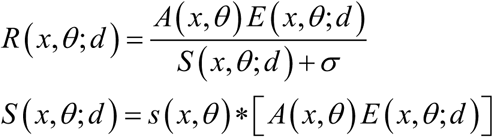

where *R(x,θ;d)* is the response of a neuron with its receptive field centered at *x* and its orientation tuning centered at *θ*, receiving probe distance (*d*) from the exogenous cue, *A(x,θ)* is the attention field, *E(x,θ;d)* is the stimulus drive of the population of neurons evoked by probe distance (*d*) from the exogenous cue, *σ* is the normalization constant, *S(x,θ;d)* is the effect of the normalizing pool and represents the excitatory drive convolved by the suppressive surround, *s(x,θ)* is the suppressive field, and *\** is convolution. Parallel to the classical contrast response function [7, 10, 76] in divisive-normalization model of attention, the population gain can be considered as a function of probe distance (D0-D4) from the exogenous cue in our study. For internet-dependents, the gradient of the population gain should be steeper for ARC than AUC conditions. Therefore, we simulated our empirical data with the divisive-normalization model of attention using custom MATLAB scripts based on the code of Reynolds and Heeger with four free parameters: the gain of attention [*A(x, u)*], separately optimized for each group and each condition, the normalization constant *σ*, the orientation tuning of AF, and a scaling parameter to linearly scale simulated values to performance (the cueing effect) (Fig. 7a).

## Acknowledgements

We acknowledge the subjects for their contribution to this study. This work was supported by National Natural Science Foundation of China (32271099) and Research Center for Brain Cognition and Human Development of Guangdong Province (2024B0303390003).

## Author contributions

Conceptualization, X.Z.; Methodology, M.M., Y.C., Y.L., Z.L., W.D., and X.Z.; Formal Analysis, M.M. and Y.C.; Investigation, M.M., Y.C., Y.L., and Z.L.; Writing – Original Draft, W.D., Y.W., and X.Z.; Writing – Review and Editing, M.M., Y.C., W.D., Y.W., and X.Z.; Supervision, X.Z.

## Data Availability

The datasets and codes for this study are available at Open Science Framework, https://osf.io/qdzr7/.

## Competing interests

The authors declare no competing interests.

## Notes

### Competing Interest Statement

The authors have declared no competing interest.

### Summary of Updates

27 smoking-dependents, 24 demographic-matched HCs of smoking-dependents, 30 internet-dependents, and 28 demographic-matched HCs of internet-dependents. revised manuscript: 30 smoking-dependents, 30 demographic-matched HCs of smoking-dependents, 30 internet-dependents, and 30 demographic-matched HCs of internet-dependents.

## References

1. Corbetta, M., & Shulman, G. L. (2002). Control of goal–directed and stimulus–driven attention in the brain. Nature reviews neuroscience, 3, 201–215. 10.1038/nrn755

2. Hegdé, J., & Felleman, D. J. (2003). How selective are V1 cells for pop–out stimuli?. Journal of Neuroscience, 23, 9968–9980. 10.1523/JNEUROSCI.23-31-09968.2003

3. Huang, L., Wang, L., Shen, W., Li, M., Wang, S., Wang, X., … & Zhang, X. (2020). A source for awareness–dependent figure–ground segregation in human prefrontal cortex. Proceedings of the National Academy of Sciences, 117, 30836–30847. 10.1073/pnas.1922832117

4. Koch, C., & Ullman, S. (1987). Shifts in selective visual attention: towards the underlying neural circuitry. In Matters of intelligence: Conceptual structures in cognitive neuroscience (pp. 115–141). Dordrecht: Springer Netherlands. 10.1007/978-94-009-3833-5_5

5. Nakayama, K. & Mackeben, M. Sustained and transient components of focal visual attention. Vision Res. 29, 1631–1647 (1989).

6. Wang, L., Huang, L., Li, M., Wang, X., Wang, S., Lin, Y., & Zhang, X. (2022). An awareness–dependent mapping of saliency in the human visual system. NeuroImage, 247, 118864. 10.1016/j.neuroimage.2021.118864

7. Wang, S., Huang, L., Chen, Q., Wang, J., Xu, S., & Zhang, X. (2021). Awareness–dependent normalization framework of visual bottom–up attention. Journal of Neuroscience, 41, 9593–9607. 10.1523/JNEUROSCI.1110-21.2021

8. Yantis, S., & Jonides, J. (1984). Abrupt visual onsets and selective attention: evidence from visual search. Journal of Experimental Psychology: Human perception and performance, 10, 601. 10.1037/0096-1523.10.5.601

9. Zhang, X., Zhaoping, L., Zhou, T., & Fang, F. (2012). Neural activities in V1 create a bottom–up saliency map. Neuron, 73, 183–192. 10.1016/j.neuron.2011.10.035

10. Zhang, X., Japee, S., Safiullah, Z., Mlynaryk, N., & Ungerleider, L. G. (2016). A normalization framework for emotional attention. PLoS biology, 14, e1002578. 10.1371/journal.pbio.1002578

11. Berridge, K. C. (2012). From prediction error to incentive salience: mesolimbic computation of reward motivation. European Journal of neuroscience, 35, 1124–1143. 10.1111/j.1460-9568.2012.07990.x

12. Field, M., & Cox, W. M. (2008). Attentional bias in addictive behaviors: a review of its development, causes, and consequences. Drug and alcohol dependence, 97, 1–20. 10.1016/j.drugalcdep.2008.03.030

13. Robbins, S. J., & Ehrman, R. N. (2004). The role of attentional bias in substance abuse. Behavioral and Cognitive Neuroscience Reviews, 3, 243–260. 10.1177/1534582305275423

14. Rooke, S. E., Hine, D. W., & Thorsteinsson, E. B. (2008). Implicit cognition and substance use: A meta–analysis. Addictive behaviors, 33, 1314–1328. 10.1016/j.addbeh.2008.06.009

15. Wilcockson, T. D. W., & Pothos, E. M. (2015). Measuring inhibitory processes for alcohol–related attentional biases: Introducing a novel attentional bias measure. Addictive behaviors, 44, 88–93. 10.1016/j.addbeh.2014.12.015

16. Wilcockson, T. D., Pothos, E. M., Osborne, A. M., & Crawford, T. J. (2021). Top–down and bottom–up attentional biases for smoking–related stimuli: comparing dependent and non–dependent smokers. Addictive behaviors, 118, 106886. 10.1016/j.addbeh.2021.106886

17. Franken, I. H., Hendriks, V. M., Stam, C. J., & Van den Brink, W. (2004). A role for dopamine in the processing of drug cues in heroin dependent patients. European Neuropsychopharmacology, 14, 503–508. 10.1016/j.euroneuro.2004.02.004

18. Franken, I. H., Booij, J., & van den Brink, W. (2005). The role of dopamine in human addiction: from reward to motivated attention. European journal of pharmacology, 526, 199–206. 10.1016/j.ejphar.2005.09.025

19. Robinson, T. E., & Berridge, K. C. (1993). The neural basis of drug craving: an incentive–sensitization theory of addiction. Brain research reviews, 18, 247–291. 10.1016/0165-0173(93)90013-P

20. Robinson, T. E., & Berridge, K. C. (2008). The incentive sensitization theory of addiction: some current issues. Philosophical Transactions of the Royal Society B: Biological Sciences, 363, 3137–3146. 10.1098/rstb.2008.0093

21. Wise, R. A., & Robble, M. A. (2020). Dopamine and addiction. Annual review of psychology, 71, 79–106. 10.1146/annurev-psych-010418-103337

22. Carpenter, K. M., Schreiber, E., Church, S., & McDowell, D. (2006). Drug Stroop performance: relationships with primary substance of use and treatment outcome in a drug–dependent outpatient sample. Addictive behaviors, 31, 174–181. 10.1016/j.addbeh.2005.04.012

23. Cox, W. M., Hogan, L. M., Kristian, M. R., & Race, J. H. (2002). Alcohol attentional bias as a predictor of alcohol abusers’ treatment outcome. Drug and alcohol dependence, 68, 237–243. 10.1016/S0376-8716(02)00219-3

24. Luijten, M., Veltman, D. J., van den Brink, W., Hester, R., Field, M., Smits, M., & Franken, I. H. (2011). Neurobiological substrate of smoking–related attentional bias. Neuroimage, 54, 2374–2381. 10.1016/j.neuroimage.2010.09.064

25. Marissen, M. A., Franken, I. H., Waters, A. J., Blanken, P., Van Den Brink, W., & Hendriks, V. M. (2006). Attentional bias predicts heroin relapse following treatment. Addiction, 101, 1306–1312. 10.1111/j.1360-0443.2006.01498.x

26. Powell, J., Dawkins, L., West, R., Powell, J., & Pickering, A. (2010). Relapse to smoking during unaided cessation: clinical, cognitive and motivational predictors. Psychopharmacology, 212, 537–549. 10.1007/s00213-010-1975-8

27. Anderson, B. A. (2016). What is abnormal about addiction–related attentional biases?. Drug and alcohol dependence, 167, 8–14. 10.1016/j.drugalcdep.2016.08.002

28. Field, M., Marhe, R., & Franken, I. H. (2014). The clinical relevance of attentional bias in substance use disorders. CNS spectrums, 19, 225–230. 10.1017/S1092852913000321

29. Anderson, B. A. (2016a). The attention habit: How reward learning shapes attentional selection. Annals of the new York Academy of Sciences, 1369, 24–39. 10.1111/nyas.12957

30. Anderson, B. A. (2021). Relating value–driven attention to psychopathology. Current opinion in psychology, 39, 48–54. 10.1016/j.copsyc.2020.07.010

31. Anderson, B. A., Faulkner, M. L., Rilee, J. J., Yantis, S., & Marvel, C. L. (2013). Attentional bias for nondrug reward is magnified in addiction. Experimental and clinical psychopharmacology, 21, 499. 10.1037/a0034575

32. Anderson, B. A. (2013). A value–driven mechanism of attentional selection. Journal of vision, 13, 7–7. 10.1167/13.3.7

33. Anderson, B. A., Laurent, P. A., & Yantis, S. (2011). Value–driven attentional capture. Proceedings of the National Academy of Sciences, 108, 10367–10371. 10.1073/pnas.1104047108

34. Kim, H., & Anderson, B. A. (2019a). Dissociable components of experience–driven attention. Current Biology, 29, 841–845. 10.1016/j.cub.2019.01.030

35. nderson, B. A., & Yantis, S. (2013). Persistence of value–driven attentional capture. Journal of Experimental Psychology: Human Perception and Performance, 39, 6. 10.1037/a0030860

36. Anderson, B. A. (2019). Neurobiology of value–driven attention. Current opinion in psychology, 29, 27–33. 10.1016/j.copsyc.2018.11.004

37. Anderson, B. A., Kuwabara, H., Wong, D. F., Gean, E. G., Rahmim, A., Brašić, J. R., … & Yantis, S. (2016). The role of dopamine in value–based attentional orienting. Current Biology, 26, 550–555. 10.1016/j.cub.2015.12.062

38. Anderson, B. A., Kuwabara, H., Wong, D. F., Roberts, J., Rahmim, A., Brašić, J. R., & Courtney, S. M. (2017). Linking dopaminergic reward signals to the development of attentional bias: A positron emission tomographic study. NeuroImage, 157, 27–33. 10.1016/j.neuroimage.2017.05.062

39. Anderson, B. A. (2017). Going for it: The economics of automaticity in perception and action. Current Directions in Psychological Science, 26, 140–145. 10.1177/0963721416686181

40. Anderson, B. A., Folk, C. L., Garrison, R., & Rogers, L. (2016). Mechanisms of habitual approach: Failure to suppress irrelevant responses evoked by previously reward–associated stimuli. Journal of Experimental Psychology: General, 145, 796. 10.1037/xge0000169

41. Kim, H., & Anderson, B. A. (2019). Neural evidence for automatic value–modulated approach behaviour. NeuroImage, 189, 150–158. 10.1016/j.neuroimage.2018.12.050

42. Field, M., Mogg, K., Zetteler, J., & Bradley, B. P. (2004). Attentional biases for alcohol cues in heavy and light social drinkers: the roles of initial orienting and maintained attention. Psychopharmacology, 176, 88–93. 10.1007/s00213-004-1855-1

43. Townshend, J., & Duka, T. (2001). Attentional bias associated with alcohol cues: differences between heavy and occasional social drinkers. Psychopharmacology, 157, 67–74. 10.1007/s002130100764

44. Field, M., Mogg, K., Mann, B., Bennett, G. A., & Bradley, B. P. (2013). Attentional biases in abstinent alcoholics and their association with craving. Psychology of Addictive Behaviors, 27, 71. 10.1037/a0029626

45. Marhe, R., Luijten, M., Van De Wetering, B. J., Smits, M., & Franken, I. H. (2013). Individual differences in anterior cingulate activation associated with attentional bias predict cocaine use after treatment. Neuropsychopharmacology, 38, 1085–1093. 10.1038/npp.2013.7

46. Waters, A. J., Shiffman, S., Bradley, B. P., & Mogg, K. (2003). Attentional shifts to smoking cues in smokers. Addiction, 98, 1409–1417. 10.1046/j.1360-0443.2003.00465.x

47. Herrmann, K., Montaser–Kouhsari, L., Carrasco, M., & Heeger, D. J. (2010). When size matters: attention affects performance by contrast or response gain. Nature neuroscience, 13, 1554–1559. 10.1111/j.1360-0443.1991.tb01809.x

48. Reynolds, J. H., & Heeger, D. J. (2009). The normalization model of attention. Neuron, 61, 168–185. 10.1016/j.neuron.2009.01.002

49. Hopf, J. M., Boehler, C. N., Luck, S. J., Tsotsos, J. K., Heinze, H. J., & Schoenfeld, M. A. (2006). Direct neurophysiological evidence for spatial suppression surrounding the focus of attention in vision. Proceedings of the National Academy of Sciences, 103, 1053–1058. 10.1073/pnas.0507746103

50. Hopf, J. M., Boehler, C. N., Schoenfeld, M. A., Heinze, H. J., & Tsotsos, J. K. (2010). The spatial profile of the focus of attention in visual search: insights from MEG recordings. Vision research, 50, 1312–1320. 10.1016/j.visres.2010.01.015

51. Huang, L., Shen, S., Sun, Y., Ou, S., Zhang, R., de Lange, F. P., & Zhang, X. (2024). Center–surround inhibition by expectation: a neuro–computational account. bioRxiv, 2024–08. 10.1101/2024.08.26.609781

52. Mangun, G. R., & Hillyard, S. A. (1988). Spatial gradients of visual attention: behavioral and electrophysiological evidence. Electroencephalography and clinical Neurophysiology, 70, 417–428. 10.1016/0013-4694(88)90019-3

53. Tkacz–Domb, S., & Yeshurun, Y. (2018). The size of the attentional window when measured by the pupillary response to light. Scientific Reports, 8, 11878. 10.1038/s41598-018-30343-7

54. Brefczynski–Lewis, J. A., Datta, R., Lewis, J. W., & DeYoe, E. A. (2009). The topography of visuospatial attention as revealed by a novel visual field mapping technique. Journal of Cognitive Neuroscience, 21, 1447–1460. 10.1162/jocn.2009.21005

55. Chapman, A. F., & Störmer, V. S. (2024). Representational structures as a unifying framework for attention. Trends in Cognitive Sciences, 28, 416–427. 10.1016/j.tics.2024.01.002

56. Shipp, S. (2004). The brain circuitry of attention. Trends in cognitive sciences, 8, 223–230. 10.1016/j.tics.2004.03.004

57. Liu, Z., Li, Y. H., Cui, Z. Y., Li, L., Nie, X. Q., Yu, C. D., … & Wang, C. (2022). Prevalence of tobacco dependence and associated factors in China: findings from nationwide China Health Literacy Survey during 2018–19. The Lancet Regional Health–Western Pacific, 24. 10.1016/j.lanwpc.2022.100464

58. Fan, T., Twayigira, M., Song, L., Luo, X., Huang, C., Gao, X., & Shen, Y. (2023). Prevalence and associated factors of internet addiction among Chinese adolescents: association with childhood trauma. Frontiers in public health, 11, 1172109. 10.3389/fpubh.2023.1172109

59. Heatherton, T. F., Kozlowski, L. T., Frecker, R. C., & FAGERSTROM, K. O. (1991). The Fagerström test for nicotine dependence: a revision of the Fagerstrom Tolerance Questionnaire. British journal of addiction, 86, 1119–1127. 10.1111/j.1360-0443.1991.tb01879.x

60. Beck, A. T., Steer, R. A., & Brown, G. (1996). Beck depression inventory–II. Psychological assessment. 10.1037/t00742-000

61. Raven, J. (2003). Raven progressive matrices. In Handbook of nonverbal assessment (pp. 223–237). Boston, MA: Springer US. 10.1007/978-1-4615-0153-4_11

62. American Psychiatric Association, D. S. M. T. F., & American Psychiatric Association, D. S. (2013). Diagnostic and statistical manual of mental disorders: DSM–5 (Vol. 5, No. 5). Washington, DC: American psychiatric association. 10.1176/appi.books.9780890425596

63. Young, K. S. (2004). Internet addiction: A new clinical phenomenon and its consequences. American behavioral scientist, 48, 402–415. 10.1177/00027642042702

64. Pessiglione, M., Seymour, B., Flandin, G., Dolan, R. J., & Frith, C. D. (2006). Dopamine–dependent prediction errors underpin reward–seeking behaviour in humans. Nature, 442, 1042–1045. 10.1038/nature05051

65. Pessiglione, M., Petrovic, P., Daunizeau, J., Palminteri, S., Dolan, R. J., & Frith, C. D. (2008). Subliminal instrumental conditioning demonstrated in the human brain. Neuron, 59, 561–567. 10.1016/j.neuron.2008.07.005

66. Palminteri, S., Justo, D., Jauffret, C., Pavlicek, B., Dauta, A., Delmaire, C., … & Pessiglione, M. (2012). Critical roles for anterior insula and dorsal striatum in punishment–based avoidance learning. Neuron, 76, 998–1009. 10.1016/j.neuron.2012.10.017

67. Palminteri, S., Khamassi, M., Joffily, M. & Coricelli, G. Contextual modulation of value signals in reward and punishment learning. Nat. Commun. 6, 8096 (2015).

68. O’Doherty, J., Dayan, P., Schultz, J., Deichmann, R., Friston, K., & Dolan, R. J. (2004). Dissociable roles of ventral and dorsal striatum in instrumental conditioning. science, 304, 452–454. 10.1126/science.1094285

69. Samejima, K., Ueda, Y., Doya, K., & Kimura, M. (2005). Representation of action–specific reward values in the striatum. Science, 310, 1337–1340. 10.1126/science.1115270

70. Sutton, R. S., & Barto, A. G. (2018). Reinforcement learning: An introduction (2end ed). MIT press. https://mitpress.mit.edu/9780262039246/reinforcement-learning

71. Shen, S., Sun, Y., Lu, J., Li, C., Chen, Q., Mo, C., … & Zhang, X. (2024). Profiles of visual perceptual learning in feature space. Iscience, 27. 10.1016/j.isci.2024.109128

72. Akaike, H. (1973). Maximum likelihood identification of Gaussian autoregressive moving average models. Biometrika, 60, 255–265. 10.1093/biomet/60.2.255

73. Schwarz, G. (1978). Estimating the dimension of a model. The annals of statistics, 461–464. http://www.jstor.org/stable/2958889

74. Burnham, K. P., & Anderson, D. R. (2002). Model selection and multimodel inference: A practical information–theoretic approach (2nd ed.). New York, NY: Springer. https://link.springer.com/book/10.1007/b97636

75. Wagenmakers, E. J. (2007). A practical solution to the pervasive problems of p values. Psychonomic bulletin & review, 14, 779–804. 10.3758/BF03194105

76. Martınez–Trujillo, J. C., & Treue, S. (2002). Attentional modulation strength in cortical area MT depends on stimulus contrast. Neuron, 35, 365–370. 10.1016/S0896-6273(02)00778-X

77. Anderson, B. A., Kronemer, S. I., Rilee, J. J., Sacktor, N., & Marvel, C. L. (2016). Reward, attention, and HIV–related risk in HIV+ individuals. Neurobiology of disease, 92, 157–165. 10.1016/j.nbd.2015.10.018

78. Connor, C. E., Gallant, J. L., Preddie, D. C., & Van Essen, D. C. (1996). Responses in area V4 depend on the spatial relationship between stimulus and attention. Journal of neurophysiology, 75, 1306–1308. 10.1152/jn.1996.75.3.1306

79. Connor, C. E., Preddie, D. C., Gallant, J. L., & Van Essen, D. C. (1997). Spatial attention effects in macaque area V4. Journal of Neuroscience, 17, 3201–3214. 10.1523/JNEUROSCI.17-09-03201.1997

80. Schall, J. D., & Hanes, D. P. (1993). Neural basis of saccade target selection in frontal eye field during visual search. Nature, 366, 467–469. 10.1038/366467a0

81. Downing, C. J. (1988). Expectancy and visual–spatial attention: effects on perceptual quality. Journal of Experimental Psychology: Human perception and performance, 14, 188. 10.1037/0096-1523.14.2.188

82. Handy, T. C., Kingstone, A., & Mangun, G. R. (1996). Spatial distribution of visual attention: Perceptual sensitivity and response latency. Perception & Psychophysics, 58, 613–627. 10.3758/BF03213094

83. Henderson, J. M., & Macquistan, A. D. (1993). The spatial distribution of attention following an exogenous cue. Perception & psychophysics, 53, 221–230. 10.3758/BF03211732

84. LaBerge, D. (1983). Spatial extent of attention to letters and words. Journal of experimental psychology: Human perception and performance, 9, 371. 10.1037/0096-1523.9.3.371

85. Posner, M. I. (2016). Orienting of attention: Then and now. Quarterly journal of experimental psychology, 69, 1864–1875. 10.1080/17470218.2014.937446

86. Shulman, G. L., Wilson, J., & Sheehy, J. B. (1985). Spatial determinants of the distribution of attention. Perception & Psychophysics, 37, 59–65. 10.3758/BF03207139

87. Couperus, J. W., & Lydic, K. O. (2019). Attentional set and the gradient of visual spatial attention. Neuroscience letters, 712, 134495. 10.1016/j.neulet.2019.134495

88. Eimer, M. (1997). Attentional selection and attentional gradients: An alternative method for studying transient visual-spatial attention. Psychophysiology, 34, 365–376. 10.1111/j.1469-8986.1997.tb02407.x

89. Moran, J., & Desimone, R. (1985). Selective attention gates visual processing in the extrastriate cortex. Science, 229, 782–784. 10.1126/science.4023713

90. Schall, J. D., Sato, T. R., Thompson, K. G., Vaughn, A. A., & Juan, C. H. (2004). Effects of search efficiency on surround suppression during visual selection in frontal eye field. Journal of Neurophysiology, 91, 2765–2769. 10.1152/jn.00780.2003

91. Bahcall, D. O., & Kowler, E. (1999). Attentional interference at small spatial separations. Vision research, 39, 71–86. 10.1016/S0042-6989(98)00090-X

92. Mounts, J. R. (2000). Evidence for suppressive mechanisms in attentional selection: Feature singletons produce inhibitory surrounds. Perception & psychophysics, 62, 969–983. 10.3758/BF03212082

93. Müller, N. G., Mollenhauer, M., Rösler, A., & Kleinschmidt, A. (2005). The attentional field has a Mexican hat distribution. Vision research, 45, 1129–1137. 10.1016/j.visres.2004.11.003

94. Boehler, C. N., Tsotsos, J. K., Schoenfeld, M. A., Heinze, H. J., & Hopf, J. M. (2009). The center–surround profile of the focus of attention arises from recurrent processing in visual cortex. Cerebral Cortex, 19, 982–991. 10.1093/cercor/bhn139

95. Boehler, C. N., Tsotsos, J. K., Schoenfeld, M. A., Heinze, H. J., & Hopf, J. M. (2011). Neural mechanisms of surround attenuation and distractor competition in visual search. Journal of Neuroscience, 31, 5213–5224. 10.1523/JNEUROSCI.6406-10.2011

96. Müller, N. G., & Kleinschmidt, A. (2004). The attentional ‘spotlight’s’ penumbra: center–surround modulation in striate cortex. Neuroreport, 15, 977–980. 10.1097/00001756-200404290-00009

97. Tsotsos, J. K., Culhane, S. M., Wai, W. Y. K., Lai, Y., Davis, N., & Nuflo, F. (1995). Modeling visual attention via selective tuning. Artificial intelligence, 78, 507–545. 10.1016/0004-3702(95)00025-9

98. Tsotsos, J. K., Rodríguez–Sánchez, A. J., Rothenstein, A. L., & Simine, E. (2008). The different stages of visual recognition need different attentional binding strategies. Brain research, 1225, 119–132. 10.1016/j.brainres.2008.05.038

99. Desimone, R., & Duncan, J. (1995). Neural mechanisms of selective visual attention. Annual review of neuroscience, 18, 193–222. 10.1146/annurev.ne.18.030195.001205

100. Yeshurun, Y. (2019). The spatial distribution of attention. Current Opinion in Psychology, 29, 76–81. 10.1016/j.copsyc.2018.12.008

101. Schallmo, M. P., Kolodny, T., Kale, A. M., Millin, R., Flevaris, A. V., Edden, R. A., … & Murray, S. O. (2020). Weaker neural suppression in autism. Nature communications, 11, 2675. 10.1038/s41467-020-16495-z

102. Robertson, C. E., Kravitz, D. J., Freyberg, J., Baron–Cohen, S., & Baker, C. I. (2013). Tunnel vision: sharper gradient of spatial attention in autism. Journal of Neuroscience, 33, 6776–6781. 10.1523/JNEUROSCI.5120-12.2013

103. Rosenberg, A., Patterson, J. S., & Angelaki, D. E. (2015). A computational perspective on autism. Proceedings of the National Academy of Sciences, 112, 9158–9165. 10.1073/pnas.151058311

104. Kaur, G., Anand, R., & Chakrabarty, M. (2023). Trait anxiety influences negative affect–modulated distribution of visuospatial attention. Neuroscience, 509, 145–156. 10.1016/j.neuroscience.2022.11.034

105. Gu, L., Yang, X., Li, L. M. W., Zhou, X., & Gao, D. G. (2017). Seeing the big picture: Broadening attention relieves sadness and depressed mood. Scandinavian Journal of Psychology, 58, 324–332. 10.1111/sjop.12376

106. Christianson, S. Å. (1992). Emotional stress and eyewitness memory: a critical review. Psychological bulletin, 112, 284. 10.1037/0033-2909.112.2.284

107. Schmitz, T. W., De Rosa, E., & Anderson, A. K. (2009). Opposing influences of affective state valence on visual cortical encoding. Journal of Neuroscience, 29, 7199–7207. 10.1523/JNEUROSCI.5387-08.2009

108. Green, C. S., & Bavelier, D. (2007). Action–video–game experience alters the spatial resolution of vision. Psychological science, 18, 88–94. 10.1111/j.1467-9280.2007.01853.x

109. Green, C. S., & Bavelier, D. (2003). Action video game modifies visual selective attention. Nature, 42, 534–537. 10.1038/nature01647

110. Shaffer, H. J., LaPlante, D. A., LaBrie, R. A., Kidman, R. C., Donato, A. N., & Stanton, M. V. (2004). Toward a syndrome model of addiction: Multiple expressions, common etiology. Harvard review of psychiatry, 12, 367–374. 10.1080/10673220490905705

111. Shaffer, H. J., Tom, M. A., Wiley, R. C., Wong, M. F., Chan, E. M., Cheng, G. L., … & Lee, M. (2018). Using the Syndrome Model of Addiction: A preliminary consideration of psychological states and traits. International Journal of Mental Health and Addiction, 16, 1373–1393. 10.1007/s11469-018-9952-2

112. Shen, W., Liu, H., Xie, X., Liu, H., & Zhou, W. (2017). Biochemical diagnosis in substance and non–substance addiction. Substance and non–substance addiction, 169–202. 10.1007/978-981-10-5562-1_9

113. Blum, K., Noble, E. P., Sheridan, P. J., Montgomery, A., Ritchie, T., Jagadeeswaran, P., … & Cohn, J. B. (1990). Allelic association of human dopamine D2 receptor gene in alcoholism. Jama, 263, 2055–2060. 10.1001/jama.1990.03440150063027

114. Kuss, D. J., Pontes, H. M., & Griffiths, M. D. (2018). Neurobiological correlates in internet gaming disorder: a systematic literature review. Frontiers in psychiatry, 9, 166. 10.3389/fpsyt.2018.00166

115. Kuss, D. J., & Griffiths, M. D. (2012). Internet gaming addiction: A systematic review of empirical research. International journal of mental health and addiction, 10, 278–296. 10.1007/s11469-011-9318-5

116. Kuss, D. J., Griffiths, M. D., Kuss, D. J., & Griffiths, M. D. (2015). Internet addiction: a real addiction?. Internet addiction in psychotherapy, 54–104. https://link.springer.com/chapter/10.1057/9781137465078_4

117. Betz, C., Mihalic, D., Pinto, M. E., & Raffa, R. B. (2000). Could a common biochemical mechanism underlie addictions?. Journal of clinical pharmacy and therapeutics, 25, 11–20. 10.1046/j.1365-2710.2000.00260.x

118. Potenza, M. N. (2006). Should addictive disorders include non-substance-related conditions?. Addiction, 101, 142–151. 10.1111/j.1360-0443.2006.01591.x

119. Petry, N. M., Rehbein, F., Gentile, D. A., Lemmens, J. S., Rumpf, H. J., Mößle, T., … & O’Brien, C. P. (2014). An international consensus for assessing internet gaming disorder using the new DSM-5 approach. Addiction, 109, 1399–1406. 10.1111/add.12457

120. Starcevic, V., & Aboujaoude, E. (2017). Internet addiction: Reappraisal of an increasingly inadequate concept. CNS spectrums, 22, 7–13. 10.1017/S1092852915000863

121. Kuss, D. J., & Lopez–Fernandez, O. (2016). Internet addiction and problematic Internet use: A systematic review of clinical research. World journal of psychiatry, 6, 143. 10.5498/wjp.v6.i1.143

122. Kardefelt–Winther, D. (2014). A conceptual and methodological critique of internet addiction research: Towards a model of compensatory internet use. Computers in human behavior, 31, 351–354. 10.1016/j.chb.2013.10.059

123. Kardefelt-Winther, D., Heeren, A., Schimmenti, A., Van Rooij, A., Maurage, P., Carras, M., … & Billieux, J. (2017). How can we conceptualize behavioural addiction without pathologizing common behaviours?. Addiction, 112, 1709–1715. 10.1111/add.13763

124. Przybylski, A. K., Weinstein, N., & Murayama, K. (2017). Internet gaming disorder: Investigating the clinical relevance of a new phenomenon. American Journal of Psychiatry, 174, 230–236. 10.1176/appi.ajp.2016.1602022

125. Davies, J. B. (1998). Pharmacology versus social process: Competing or complementary views on the nature of addiction?. Pharmacology & therapeutics, 80, 265–275. 10.1016/S0163-7258(98)00031-X

126. Li, R., Polat, U., Makous, W., & Bavelier, D. (2009). Enhancing the contrast sensitivity function through action video game training. Nature neuroscience, 12, 549–551. 10.1038/nn.2296

127. Gil–Gómez, J. A., Lloréns, R., Alcañiz, M., & Colomer, C. (2011). Effectiveness of a Wii balance board–based system (eBaViR) for balance rehabilitation: a pilot randomized clinical trial in patients with acquired brain injury. Journal of neuroengineering and rehabilitation, 8, 1–10. 10.1186/1743-0003-8-30

128. Dunn, E. W., Gilbert, D. T., & Wilson, T. D. (2011). If money doesn’t make you happy, then you probably aren’t spending it right. Journal of consumer psychology, 21, 115–125. 10.1016/j.jcps.2011.02.002

129. Ngetich, R., Burleigh, T. L., Czakó, A., Vékony, T., Németh, D., & Demetrovics, Z. (2023). Working memory performance in disordered gambling and gaming: A systematic review. Comprehensive Psychiatry, 152408. 10.1016/j.comppsych.2023.152408

130. Spence, I., & Feng, J. (2010). Video games and spatial cognition. Review of general psychology, 14, 92–104. 10.1037/a0019491

131. Greitemeyer, T., & Osswald, S. (2011). Playing prosocial video games increases the accessibility of prosocial thoughts. The Journal of social psychology, 151, 121–128. 10.1080/00224540903365588

132. Barlett, C. P., Vowels, C. L., Shanteau, J., Crow, J., & Miller, T. (2009). The effect of violent and non–violent computer games on cognitive performance. Computers in Human Behavior, 25, 96–102. 10.1016/j.chb.2008.07.008

133. Martin, P. R., & Petry, N. M. (2005). Are non–substance–related addictions really addictions?. American Journal on Addictions, 14, 1–7. 10.1080/10550490590899808

134. Henningfield, J. E., Cohen, C., & Slade, J. D. (1991). Is nicotine more addictive than cocaine?. British journal of addiction, 86, 565–569.

135. Kozlowski, L. T., Wilkinson, D. A., Skinner, W., Kent, C., Franklin, T., & Pope, M. (1989). Comparing Tobacco Cigarette Dependence With Other Drug Dependencies: Greater or Equal’Difficulty Quitting’and’Urges to Use,’but Less’ Pleasure’From Cigarettes. Jama, 261, 898–901 10.1001/jama.1989.03420060114043

136. Hughes, J. R. (2001). Why does smoking so often produce dependence? A somewhat different view. Tobacco Control, 10, 62–64. 10.1136/tc.10.1.62

137. Warburton, D. M., Revell, A., & Walters, A. C. (1988). Nicotine as a resource. In M. J. Rand & K. Thurau (Eds.), Pharmacology of nicotine (pp. 359–373). London: ICSU Press. 10.1136/tc.10.1.62

138. Balfour, D. J. (2004). The neurobiology of tobacco dependence: a preclinical perspective on the role of the dopamine projections to the nucleus. Nicotine & Tobacco Research, 6, 899–912. 10.1080/14622200412331324965

139. Hughes, J. R. (2006). Should criteria for drug dependence differ across drugs?. Addiction, 101, 134–141. 10.1111/j.1360-0443.2006.01588.x

140. Frenk, H., & Dar, R. (2000). A critique of nicotine addiction. Norwell, MA: Kluwer Academic Publishers. https://link.springer.com/book/10.1007/b111440

141. Atrens, D. M. (2001). Nicotine as an addictive substance: a critical examination of the basic concepts and empirical evidence. Journal of Drug Issues, 31, 325–394. 10.1177/002204260103100202

