## Supplemental information for "Attention field as a cognitive-behavioral marker for demarcating internet- but not smoking-addiction from reward"

**Supplemental information**
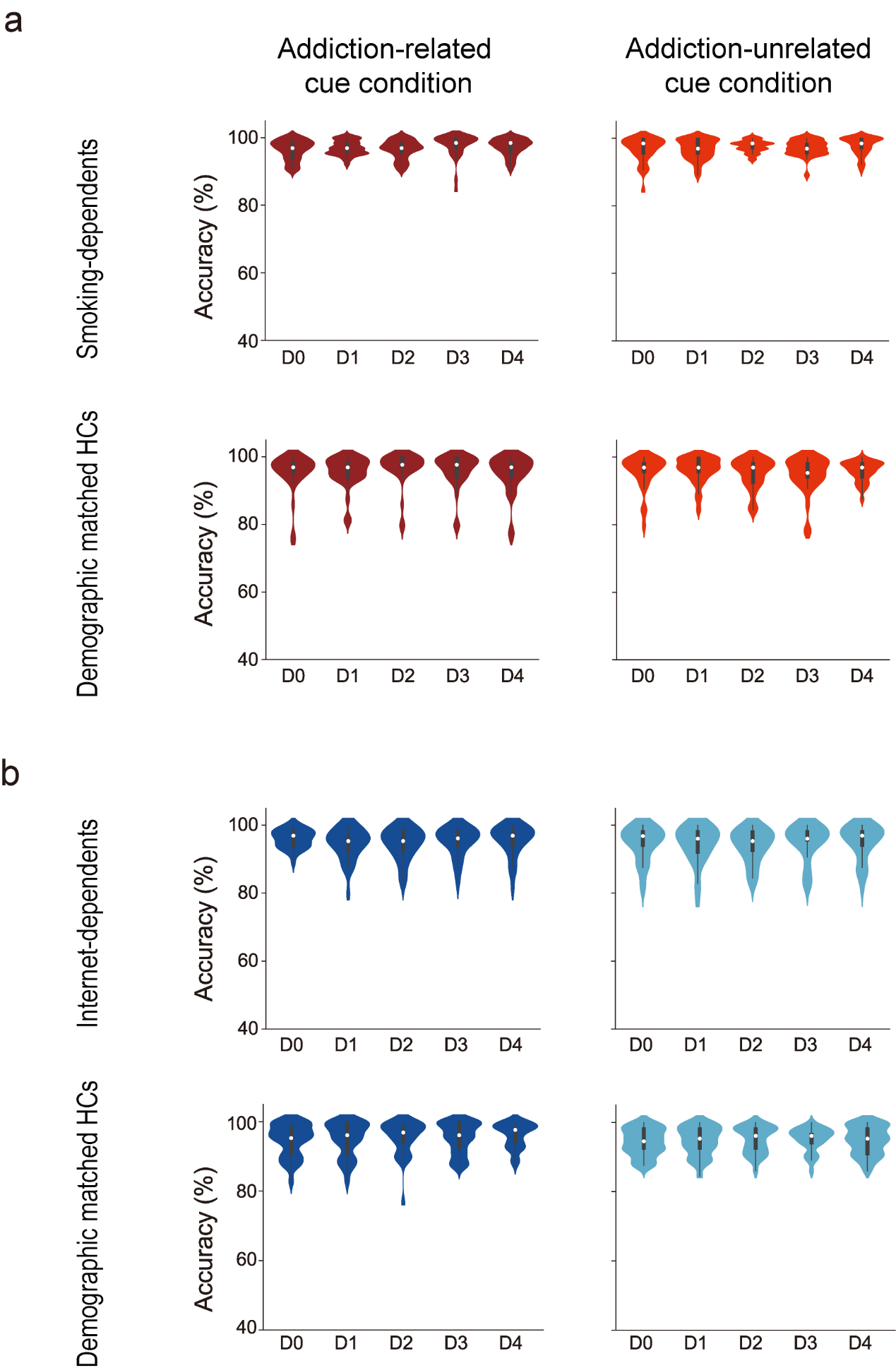

### Fig. S1. Accuracy of each distance during each group and each condition in the Test phase. For smoking-dependents and their HCs (a), their accuracies of conditions were submitted to a three-factor mixed ANOVA with group (addictions and HCs) as the between-subjects factor, and with cue (addiction-related and addiction-unrelated) and distance (D0-D4) as within-subjects factors. Neither the main effects (all *P* > 0.106, η_p_^2^ < 0.044) nor interactions (all *P* > 0.097, η_p_^2^ < 0.044) were significant, suggesting no difference in accuracies across conditions. Similarly, for internet-dependents and their HCs (b), their accuracies of conditions were also submitted to a three-factor mixed ANOVA with group (addictions and HCs) as the between-subjects factor, and with cue (addiction-related and addiction-unrelated) and distance (D0-D4) as within-subjects factors. Neither the main effects (all *P* > 0.169, η_p_^2^ < 0.032) nor interactions (all *P* > 0.268, η_p_^2^ < 0.021) were significant, confirming no difference in accuracies across conditions. Smoothed density plot denotes the distribution of the sample, box plot denotes the quantile and median scores, and error bars denote 1 SEM calculated across subjects.

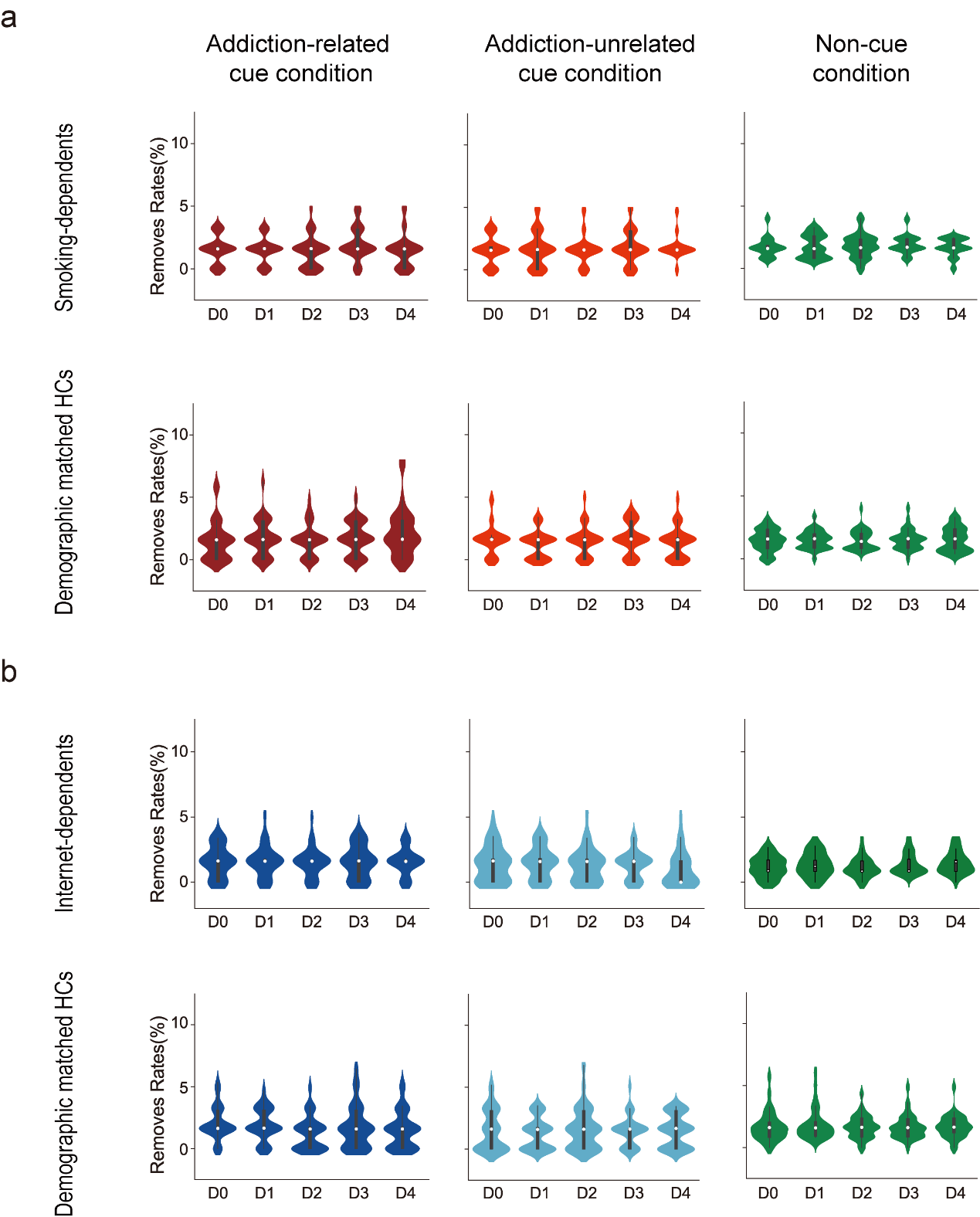

### Fig. S2. Removal rate of each distance during each group and each condition in the Test phase. Removal rates (i.e., correct reaction times shorter than 200 ms and beyond three standard deviations from the mean reaction time in each condition were removed) of smoking-dependents and their HCs (a), and of internet-dependents and their HCs (b). Similar to the accuracy in Fig. S1, these removal rates were submitted to a three-factor mixed ANOVA with group (addictions and HCs) as the between-subjects factor, and with cue (addiction-related and addiction-unrelated) and distance (D0-D4) as within-subjects factors. Neither the main effects (Smoking-dependents and their HCs: all *P* > 0.425, η_p_^2^ < 0.011; Internet-dependents and their HCs: all *P* > 0.110, η_p_^2^ < 0.044) nor interactions (Smoking-dependents and their HCs: all *P* > 0.225, η_p_^2^ < 0.021; Internet-dependents and their HCs: all *P >* 0.115, η_p_^2^ < 0.042) were significant, indicating no difference in removal rates across conditions. Smoothed density plot denotes the distribution of the sample, box plot denotes the quantile and median scores, and error bars denote 1 SEM calculated across subjects.

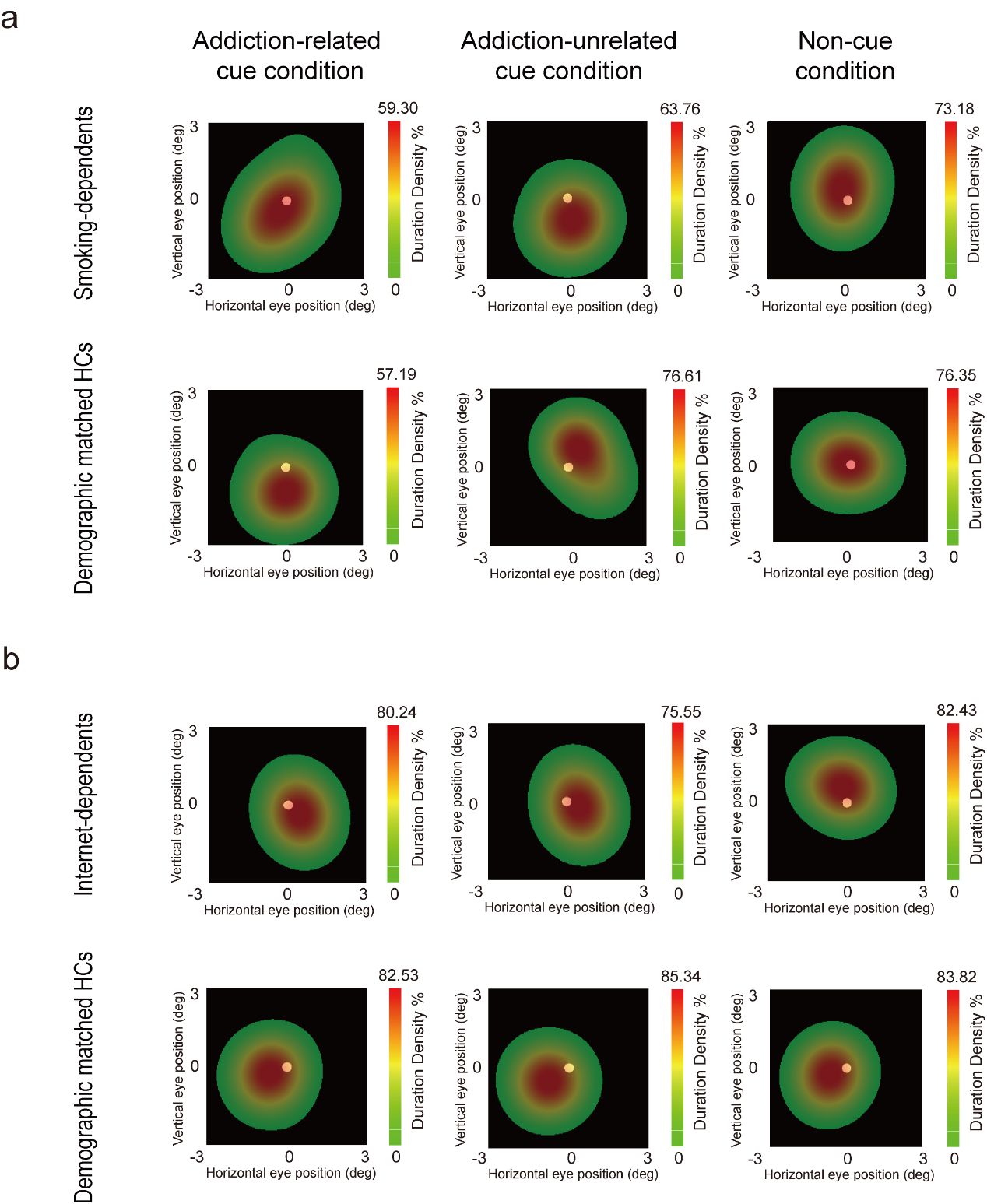

**Fig. S3. Eye movement data in the Test phase.** Horizontal and vertical eye positions after removing blinks and artifacts of the addiction-related cue (left), addiction-unrelated cue (middle), and non-cue (right) conditions for smoking-dependents and their HCs (**a**), and for internet-dependents and their HCs (**b**). Subjects’ eye movements were small (< 3.0°) and not systematically different among conditions (all p > 0.05).

**Table S1. Results of covariate analyses with demographic characteristics**

| Smoking-dependents & HCs | | | | | | | | | |
| --- | --- | --- | --- | --- | --- | --- | --- | --- | --- |
|  | Attentional effect (Peak) | | | Attentional effect (Mean) | | | FWHM bandwidth | | |
|  | Main effect (Group) | Main effect (Cue) | Interaction  between two | Main effect (Group) | Main effect (Cue) | Interaction  between two | Main effect (Group) | Main effect (Cue) | Interaction  between two |
| Age | 0.612 | 0.489 | 0.741 | 0.695 | 0.176 | 0.924 | 0.674 | 0.429 | 0.876 |
| Depression | 0.627 | 0.212 | 0.476 | 0.664 | 0.253 | 0.598 | 0.864 | 0.338 | 0.955 |
| IQ | 0.523 | 0.820 | 0.603 | 0.536 | 0.263 | 0.714 | 0.620 | 0.414 | 0.890 |
| IGD | 0.680 | 0.205 | 0.540 | 0.727 | 0.097 | 0.546 | 0.556 | 0.440 | 0.980 |
| YIAT | 0.679 | 0.147 | 0.485 | 0.724 | 0.163 | 0.564 | 0.978 | 0.628 | 0.978 |
| Internet-dependents & HCs | | | | | | | | | |
|  | Attentional effect (Peak) | | | Attentional effect (Mean) | | | FWHM bandwidth | | |
|  | Main effect (Group) | Main effect (Cue) | Interaction  between two | Main effect (Group) | Main effect (Cue) | Interaction  between two | Main effect (Group) | Main effect (Cue) | Interaction  between two |
| Age | 0.943 | 0.831 | 0.184 | 0.391 | 0.933 | 0.092 | 0.609 | 0.393 | **0.018*** |
| Depression | 0.838 | 0.120 | 0.144 | 0.347 | 0.358 | 0.084 | 0.506 | **0.024*** | **0.024*** |
| IQ | 0.751 | 0.923 | 0.169 | 0.301 | 0.977 | 0.092 | 0.620 | 0.639 | **0.021*** |

Note that, for internet-dependents and their HCs, the interactions between group and cue were all significant with the age (F(1, 57) = 5.971, *p* = 0.018, η_p_^2^ = 0.095), depression (F(1, 57) = 5.364, *p* = 0.024, η_p_^2^ = 0.086), or IQ (F(1, 57) = 5.611, *p* = 0.021, η_p_^2^ = 0.090) as the covariate. Further simple effect analysis showed that, the FWHM bandwidth of ARC condition was significantly smaller than that of AUC condition for internet-dependents (Age as the covariate: F(1, 28) = 0.462, *p* = 0.502, η_p_^2^ = 0.016; Depression as the covariate: F(1, 28) = 7.032, *p* = 0.013, η_p_^2^ = 0.201; IQ as the covariate: F(1, 28) = 0.302, *p* = 0.587, η_p_^2^ = 0.11) , but not for their HCs (Age as the covariate: F(1, 28) = 0.278, *p* = 0.602, η_p_^2^ = 0.010; Depression as the covariate: F(1, 28) = 0.453, *p* = 0.506, η_p_^2^ = 0.016; IQ as the covariate: F(1, 28) = 0.007, *p* = 0.933, η_p_^2^ < 0.001); the FWHM bandwidth of internet-dependents was significantly smaller than that of HCs for ARC condition (Age as the covariate: F(1, 57) = 4.808, *p* = 0.032, η_p_^2^ = 0.078; Depression as the covariate: F(1, 57) = 5.075, *p* = 0.028, η_p_^2^ = 0.082; IQ as the covariate: F(1, 57) = 4.602, *p* = 0.036, η_p_^2^ = 0.075), but not for AUC condition (Age as the covariate: F(1, 57) = 0.778, *p* = 0.381, η_p_^2^ = 0.013; Depression as the covariate: F(1, 57) = 0.499, *p* = 0.483, η_p_^2^ = 0.009; IQ as the covariate: F(1, 57) = 0.747, *p* = 0.391, η_p_^2^ = 0.013). These results further confirm a reduction in the AF of bottom-up attention in internet-dependents.
